# Three-dimensional genome structure and chromatin accessibility reorganization during *in vivo* induction of human T cell tolerance

**DOI:** 10.1101/2020.03.11.988253

**Authors:** Ruifeng Li, Huidong Guo, Yingping Hou, Shuoshuo Liu, Ming Wang, Ting Peng, Xiang-Yu Zhao, Liming Lu, Yali Han, Yiming Shao, Ying-Jun Chang, Cheng Li, Xiao-Jun Huang

## Abstract

Achieving T cell tolerance is a pivotal goal for the field of transplantation and autoimmune diseases. Here, we characterized the gene expression profiles, 3D genome architecture and chromatin accessibility in human steady-state and tolerant T cells, which had been induced in healthy donors by granulocyte-colony-stimulating factor *in vivo*. We provided the first high-resolution 3D genomic landscape of human tolerant T cells *in vivo* and identified highly expressed suppressor of cytokine signaling 1 (*SOCS1*), which is essential for maintaining T cell tolerance and was validated by *ex vivo* experiments. Mechanistically, SOCS1 is activated by STAT3, which mediates a new interaction between the *SOCS1* locus and downstream super-enhancers and is accompanied by the disruption of the CTCF loop between the *SOCS1* locus and upstream heterochromatin. This competitive regulation pattern between STAT3 and CTCF is present in the whole genome. Our study defines a regulatory model of transcription factors and provides insight into the induction of immune tolerance.

## INTRODUCTION

T cells, including helper cells (CD3^+^CD4^+^) and cytotoxic T-cells (CD3^+^CD8^+^), are major cell subsets in the immune system (Janssen et al., 2003). The activation and tolerance of T cells are tightly regulated to ensure effective elimination of foreign antigens while maintaining tolerance to self-antigens (Handel et al., 2018; Jenkins and Schwartz, 1987; Wu and Reddy, 2017). T cell tolerance could be induced by several strategies, such as blockade of costimulatory signals, transforming growth factor beta and regulatory T cells (Treg) (Liu et al., 2019; McCarron et al., 2019; Soto-Nieves et al., 2009; Xu et al., 2018), to prevent autoimmune diseases or to allow successful allogeneic stem cell transplantation (Allo-SCT) and allogeneic organ transplantation (Handel et al., 2018; Wu and Reddy, 2017), both of which have major clinical implications.

Previous studies on the induction of T cell tolerance have mainly focused on single genes or transcription factors (TFs), such as Egr-2, nuclear factor of activated T cells (NFAT), and Jun (Lynn et al., 2019; Safford et al., 2005; Soto-Nieves et al., 2009). Recent advances in genome sequencing technologies, including Hi-C (Genome-wide chromosome conformation capture), assays for transposase-accessible chromatin with high-throughput sequencing (ATAC-seq) and RNA-seq, have enabled gene expression and epigenetic measurements and have revealed variability in immune-cell development and aging (Calderon et al., 2019; Miraldi et al., 2019; Philip et al., 2017). For example, by RNA-seq and ATAC-seq, Mognol et al. defined a pattern of chromatin accessibility specific for T-cell exhaustion (Mognol et al., 2017), which was characterized by enrichment for consensus binding motifs for the NR4A and NFAT TFs. More recently, the transcription factor NR4A1 was identified as a key mediator of T cell dysfunction based on a genome-wide transcriptomic assessment of mouse tolerant T cells (Liu et al., 2019). However, the multiomic landscapes of human tolerant T cells have not been demonstrated. Advances in multiomic technologies have led to the development of possible approaches to investigate the 3D genome architecture and chromatin accessibility in human tolerant T cells (Lieberman-Aiden et al., 2009; Nagano et al., 2015; Rao et al., 2014; Satpathy et al., 2018).

Previous studies have shown that granulocyte-colony-stimulating factor (G-CSF), initially identified as a growth factor for neutrophils, has been widely used as a mobilizer for stem/progenitor cells in allo-HSCT settings. In the past two decades, increasing evidence has supported the critical role of recombinant human G-CSF in the induction of T cell tolerance, which is characterized by decreased proliferation and interleukin-2 production, of healthy allo-SCT donors (Jun et al., 2004; MacDonald et al., 2014; Pan et al., 1995; Rutella et al., 2005). *Ex vivo* experiments also suggested that G-CSF is a strong immune regulator of T cells and directly modulates T-cell immune responses via its receptor on T cells (Franzke et al., 2003). *In vivo* experiments showed that donor T cell alloreactivity is directly modulated through binding to the G-CSF receptor expressed in T cells (MacDonald et al., 2014). These data indicate that human tolerant T cells induced by G-CSF provide a platform to elucidate the genome landscape of tolerance.

In this study, we combined multiomic technologies, including Hi-C (Lieberman-Aiden et al., 2009; Rao et al., 2014), ATAC-seq (Buenrostro et al., 2013), RNA-seq, ChIP-seq (Johnson et al., 2007) and CUT&Tag (Kaya-Okur et al., 2019), and a T cell tolerance model established after treating healthy donors with G-CSF (Jun et al., 2004; MacDonald et al., 2014; Pan et al., 1995; Rutella et al., 2005) to investigate both the transcriptional and 3D genome landscape associated with human T cell tolerance. We identified SOCS1 as a key mediator that promotes T cell tolerance and validated the results using *in vitro* experiments.

## RESULTS

### Landscape determined from multiomic analyses of human CD4^+^ T cells and CD8^+^ T cells

We assessed the genome landscape and chromatin accessibility of steady-state CD4^+^ and CD8^+^ T cells (CD4^+^ T_ss_ and CD8^+^ T_ss_, respectively) in human bone marrow (BM) (Figure 1A and Figure S1A-C). We found that CD4^+^ T_ss_ and CD8^+^ T_ss_ were distinct from each other, as shown in the dendrogram using transcriptome data (Figure 1B) and chromatin accessibility data (Figure S1D). There were 128 genes overexpressed in CD4^+^ T_ss_ and 207 genes overexpressed in CD8^+^ T_ss_. The genes of CD4, interleukin 2 receptor subunit alpha (IL-2RA), and OX40 (also called tumor necrosis factor receptor superfamily member 4, TNFSF4) are specifically expressed in the CD4^+^ T_ss_ cells, whereas CD8A, CD8B, and KLRD1 are expressed in the CD8^+^ T_ss_ cells (Figure 1C and 1D). These observations confirmed a difference in gene expression between the CD4^+^ lineage and CD8^+^ lineage (Kioussis and Ellmeier, 2002; Mingueneau et al., 2013; Monaco et al., 2019; Satpathy et al., 2018; Szabo et al., 2019). The TF Foxp3, a key regulatory gene for the development of Tregs, is highly expressed in CD4^+^ T_ss_ (Figure 1C) (Hori et al., 2003). Gene pathway enrichment analysis showed that highly expressed genes enriched in various pathways are related to the positive regulation of lymphocyte proliferation in CD4^+^ T_ss_ and graft-versus-host disease in CD8^+^ T_ss_ (Figure 1E) (Zhao et al., 2011).

**Figure 1.**
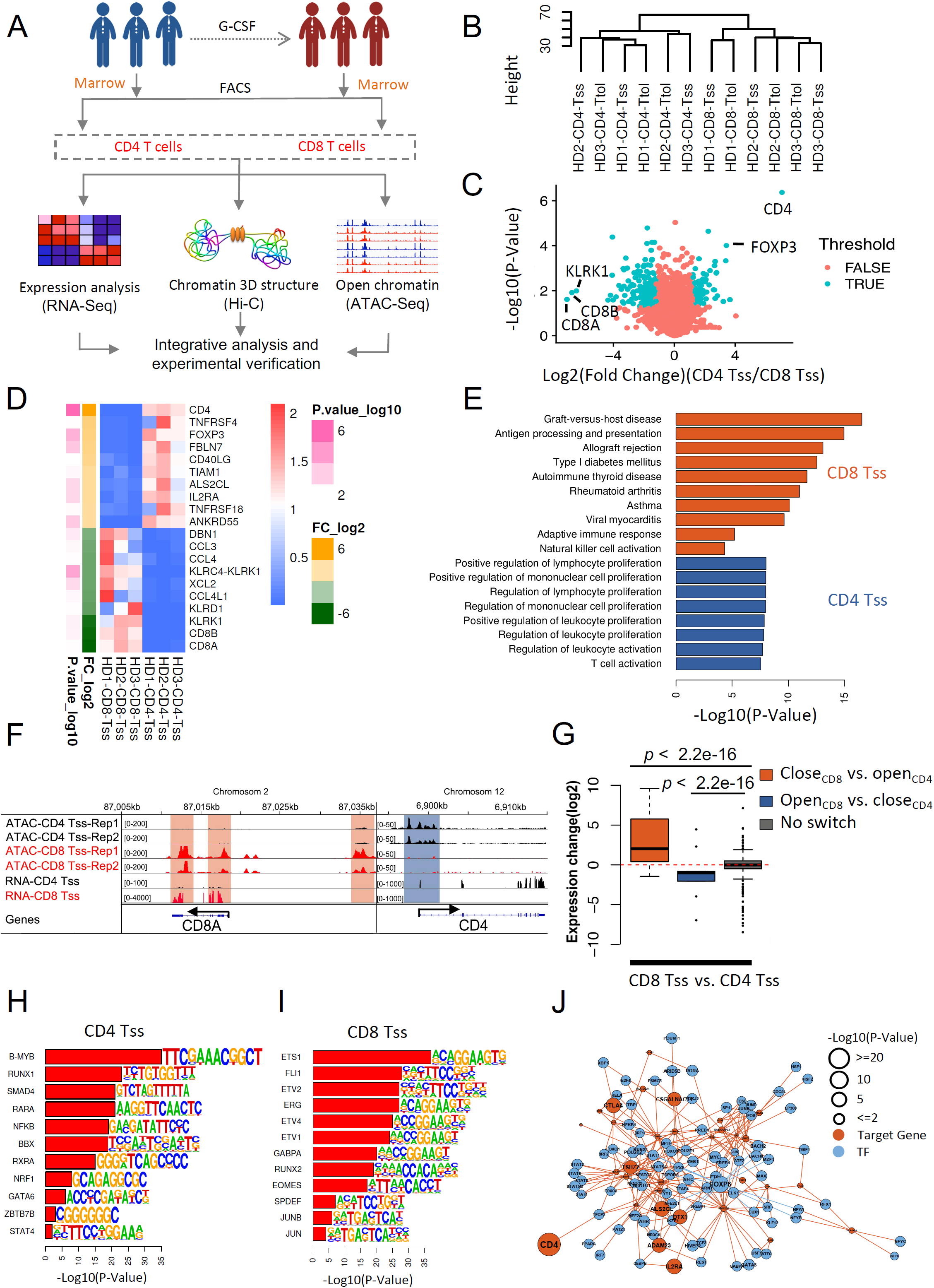
Specific genes and TFs in CD4 and CD8 cell types. **(A)** Outline of the experiments and analyses in this study. We performed in situ Hi-C, RNA-seq and ATAC-seq experiments for paired T cells (CD4 and CD8) from three healthy donors before and after G-CSF mobilization *in vivo*. **(B)** Unsupervised hierarchical clustering of the transcriptome (the first 1000 genes with the largest variance). **(C)** Volcano plot comparing CD4 T_ss_ and CD8 T_ss_. The X-axis is the fold change (log2) of CD4/CD8. There were 128 genes overexpressed in CD4 T_ss_ cells and 207 genes overexpressed in CD8 T_ss_ cells. **(D)** Corresponding to C, the top ten genes with high and low expression were identified (red: high expression; blue: low expression). **(E)** Gene pathway enrichment analysis for the highly expressed genes in CD8 T_ss_ (red) or CD4 T_ss_ (blue). **(F)** The UCSC browser views showing ATAC-seq and RNA-seq of representative genes (CD8A and CD4) in CD8 and CD4 cells of donor HD1. The data between different repeats show satisfactory consistency. The CD8A gene is overexpressed in CD8 cells, with higher chromatin accessibility of the promoter region of CD8A. The CD4 gene is overexpressed in CD4 cells, with higher chromatin accessibility of the promoter region of CD4. Orange represents chromatin regions with higher accessibility in CD8 cells than in CD4 cells. Blue represents the opposite situation. **(G)** Boxplots of expression changes of genes grouped by their chromatin accessibility signal changes. In the open and closed transition areas, we selected the top 100 regions with the most significant variation (CD8-specific genes: 40, CD4-specific genes: 52). Genome-wide, genes in high chromatin accessibility regions have higher RNA expression levels. Gray represents the chromatin regions with no significant difference in accessibility between CD4 T_ss_ and CD8 T_ss_. **(H)** Transcription factors predicted to have high chromatin accessibility of CD4 T_ss_ cells using ATAC-seq data by HOMER software. **(I)** Transcription factors predicted for high chromatin accessibility of CD8 T_ss_ cells using ATAC-seq data by HOMER software. **(J)** The regulatory network of overexpressed genes and active transcription factors in CD4. Blue dots represent transcription factors, and red dots represent target genes. The regulatory relationship between transcription factors and genes comes from Yan et al. (Yan et al., 2012).

Using ATAC-seq technology (Buenrostro et al., 2013), we observed higher chromatin accessibility at the promoter and upstream elements of the cell-type specific genes than that at other regions in CD4^+^ and CD8^+^ T_ss_ (Figure 1F and Figure S1E-G), consistent with the characteristics of these two cell lineages (Huang et al., 2006; Kioussis and Ellmeier, 2002). We selected the top 100 chromatin regions specifically accessible in CD4^+^ and CD8^+^ T_ss_ (40 genes in the CD8-specific accessible region and 52 genes in the CD4-specific accessible region). Genome-wide, genes in high chromatin accessibility regions had higher RNA expression levels than those in other regions (Figure 1G). We further showed the associations of active TFs, such as RUNX1, SMAD4 and ZBTB7B in CD4^+^ T_ss_, whereas EOMES and JUN in CD8^+^ T_ss_ (Figure 1H and 1I)(Mingueneau et al., 2013). There are essential TFs in the regulatory network of CD4^+^ T_ss_, such as GATA3/STAT6 and Foxp3/STAT5 (Figure 1J), similar to those of helper T cell lineages in peripheral blood (Stritesky et al., 2011; Zhu and Paul, 2010). These results highlight the feasibility and reliability of RNA-seq and ATAC-seq in investigating the genome landscape and chromatin accessibility of human T cells.

### High-resolution maps of 3D genome structures in human CD4^+^ T cells and CD8^+^ T cells

We used Hi-C technology to investigate the specific chromatin structures of human BM CD4^+^ and CD8^+^ T_ss_ (Figure 2) (Rao et al., 2014). We got high-resolution maps (10 Kb) of 3D genome structures including A/B compartments, TAD structure and loop structure of all the samples (Figure 2A). Intrachromosome interactions are more frequent than interchromosomal interactions in CD8^+^ T_ss_ (cis interaction ratio: 42%-56%, Figure S1A). HiCRep’s SCC scores (Yang et al., 2017) of the Hi-C matrices showed that cells of different lineages have different 3D genome structures (Figure 2B). The loop length of CD4^+^ T_ss_ was significantly (median length, CD4 T_ss_: 210 Kb, CD8 T_ss_ 190 Kb) longer than that of CD8^+^ T_ss_ (Figure 2C and 2D). This finding indicates that the chromatin structure of CD4^+^ T_ss_ is more variable than that of CD8^+^ T_ss_ at the level of loops. Separately examining the matrices of CD4^+^ T_ss_ and CD8^+^ T_ss_, we found 9,787 loops in the former and 7,481 loops in the latter, with 5,239 in both lists. The accuracy of the resulting loop calls was supported by high-scoring aggregate peak analysis (APA) plots (Rao et al., 2014) (Figure 2E). The loops, associated with CTCF and YY1, which have been proven to have universal chromosomal architecture functions in humans (Rao et al., 2014; Weintraub et al., 2017), are similar between CD4^+^ T_ss_ and CD8^+^ T_ss_ (Figure 2F). Some loops, associated with PRDM1 and ZEB1, are specific for CD4 T_ss_, while others, including IRF4 and BBX, are specific for CD8 T_ss_ (Figure 2F). More than 55% of the up-regulated genes are located in the loop anchor region, which indicates that the regulation of these genes may be related to the 3D genome structure (Figure 2G). The gene expression of Foxp3, IL-2RA, CD4, and TIAM1 in the loop anchor region of CD4 T_ss_ as well as the association of chromatin accessibility and 3D genome structures was detected. We showed the association of higher expression of TIAM1 as well as CD8A and CD8B with chromatin accessibility and the long-distance regulated enhancer in CD4^+^ T_ss_ and CD8^+^ T_ss_, respectively (Figure 2H and 2I). These results suggest the feasibility and reliability of Hi-C in exploring the genomic landscape of T cells.

**Figure 2.**
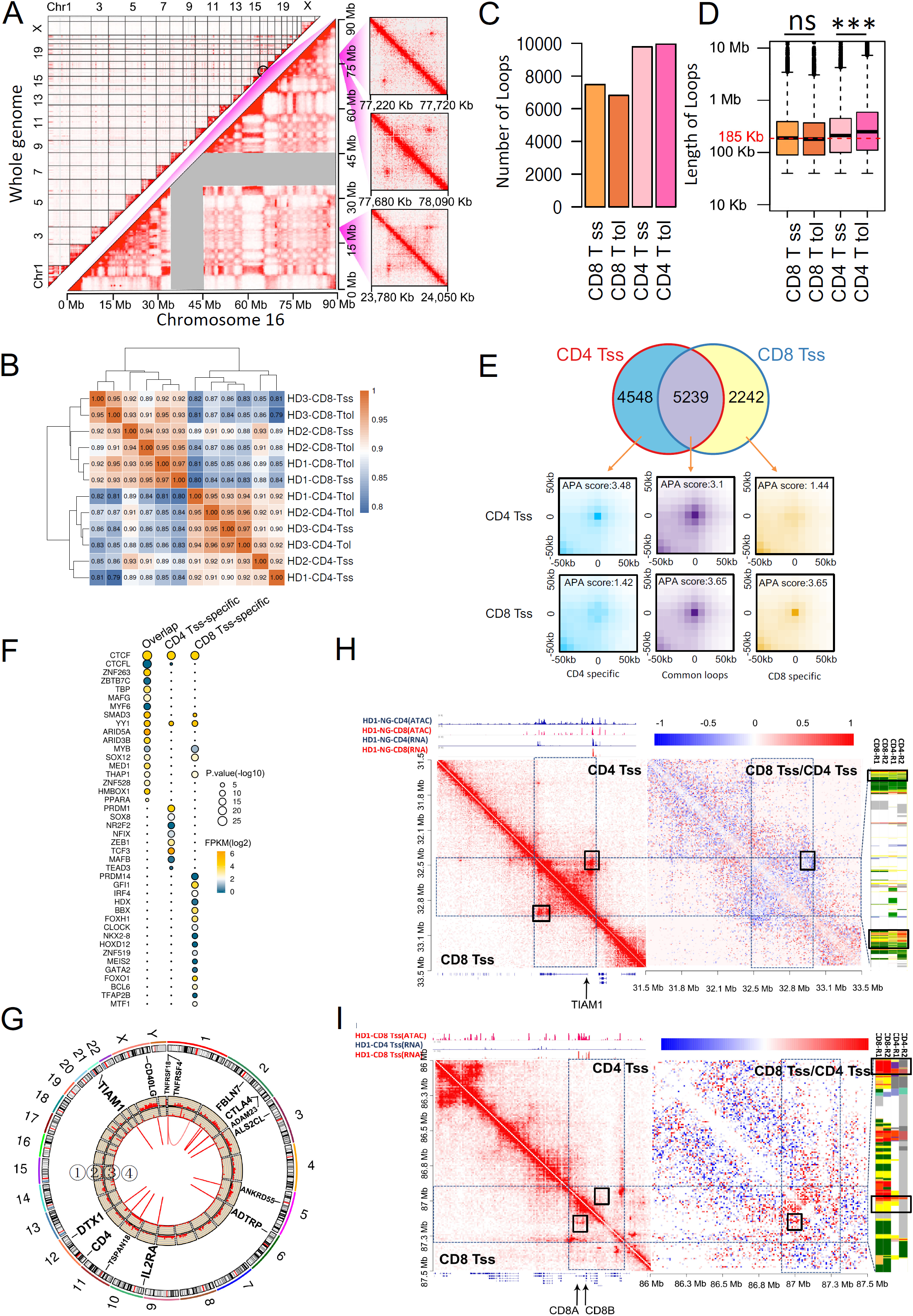
Specific chromatin structures in CD4 and CD8 cell types. **(A)** High-resolution maps of 3D genome structures. Top left: Whole genome Hi-C interaction matrix of the CD8 T_ss_ cells. Bottom right: Hi-C interaction matrix of chromosome 16 of the CD8 T_ss_ cells. Right: Examples of three loop structures. **(B)** Correlations of Hi-C matrices in all samples using HiCRep’s SCC scores (Yang et al., 2017). **(C)** The number of loops. **(D)** The length of the loops. Consistent with previous reports, the median loop length was 185 Kb (red dotted line), and the loop length of the CD4 cells increased significantly (median length, CD8 T_ss_ cells: 190 Kb, CD8 T_tol_ cells: 180 Kb, CD4 T_ss_ cells: 210 Kb, CD4 T_tol_ cells: 340 Kb). **(E)** Venn diagram of the chromatin loops of the CD4 T_ss_ cells and the CD8 T_ss_ cells. APA analysis was performed on the three types of loops in the Venn diagram to verify the reliability of each type of loop. **(F)** Motif results predicted for different types of loop anchors by HOMER software. **(G)** CD4 T_ss_ cells showing specific highly expressed genes in the loop anchor region. From the outer circle to the inner circle: ⍰: gene name; ⍰: gene expression levels from RNA-seq data in CD4^+^ T_ss_; ⍰: chromosome accessibility from ATAC-seq data in CD4^+^ T_ss_; ⍰ red lines: chromatin loops whose loop anchors overlap with CD4 specific genes. **(H)** This heatmap shows the representative gene TIAN1 with higher expression in CD4 T_ss_ cells than CD8 T_ss_ cells located in loops with higher interactions on chr21 (chr21:31,500,000–35,500,000) in CD4 cells than in CD8 T_ss_ cells. The heatmap on the left represents the chromosome interaction matrix (resolution is 10 kb). The upper right is the matrix of CD4 T_ss_, and the lower left is the matrix of CD8 T_ss_. The histogram above the heatmap is the chromatin accessibility and gene expression in this region. The annotation below the heatmap is the gene name. The heatmap on the right represents the log ratio of CD8 T_ss_ and CD4 T_ss_ chromatin interactions, blue represents a stronger interaction in CD4 T_ss_ cells, and red represents a stronger interaction in CD8 T_ss_ cells. The loop where the TIAM1 gene is located is blue, indicating that the interaction of the loop in CD4 T cells is significantly stronger than that of CD8 T_ss_ cells. The right side of the heatmap is the annotation of the chromHMM of the loop sitting area. Yellow represents an enhancer, red represents a promoter, and gray and white represent heterochromatin. **(I)** Similar to F, the heatmap shows representative genes CD8A and CD8B with higher expression in CD4 T_ss_ cells located in loops of higher interactions on chr2 (chr2:86,000,000–87,500,000) in CD8 T_ss_ cells than in CD4 T_ss_ cells.

### Landscape of the 3D genomic structure and chromatin accessibility in tolerant CD4^+^ and CD8^+^ T cells

T cell tolerance could be induced by *in vivo* application of G-CSF in both mice and humans (Jun et al., 2004; MacDonald et al., 2014; Pan et al., 1995; Rutella et al., 2005). Therefore, we explored the 3D genome structure and chromatin accessibility reorganization during the induction of human tolerant CD4^+^ and CD8^+^ T cells (CD4^+^ T_tol_ and CD8^+^ T_tol_) induced by G-CSF (Jun et al., 2004; MacDonald et al., 2014; Pan et al., 1995; Rutella et al., 2005). We found that CD8^+^ T_tol_ cells differentially expressed multiple genes, such as SOCS1, TNFSF8 and SEMA7A (Figure 3A-B), compared to CD8^+^ T_ss_ cells. CD8 T_tol_ showed a significant downregulation of genes related to cell activation (Figure 3C). We also found that CD8^+^ T_tol_ differentially activated TFs, such as STAT3 (Figure 3D-E), compared to CD8^+^ T_ss_. Differential expression of genes and differentially activated TFs, including TXNIP and RUNX1, between CD4^+^ T_tol_ and CD4^+^ T_ss_ were observed (Figure S2A-D). We observed a core set of gene and TF changes, including those of suppressor of cytokine signaling 1 (SOCS1), Kruppel like factor 9 (KLF9), and Fra2, in a similar manner in CD4^+^ T_tol_ and CD8^+^ T_tol_ (Figure 3A-D, Figure S2B-D, and S3A-B). However, both CD4^+^ T_tol_ and CD8^+^ T_tol_ also had changes in specific sets of genes and TFs. Notably, CD8^+^ T cell-based changes in gene and TF included that of SEMA7A and B-MYB, whereas CD4^+^ T cell-based changes included that of IRF4 and nuclear factor-κB inhibitor alpha (NFKBIA) (Figure 3B-E, Figure S2B-E). Thus, although there is a clear transcriptomic change of tolerant T cells shared by both CD4 and CD8 lineages, there were also unique differentially expressed genes and TFs in CD4^+^ and CD8^+^ T cells during tolerance induction *in vivo*.

**Figure 3.**
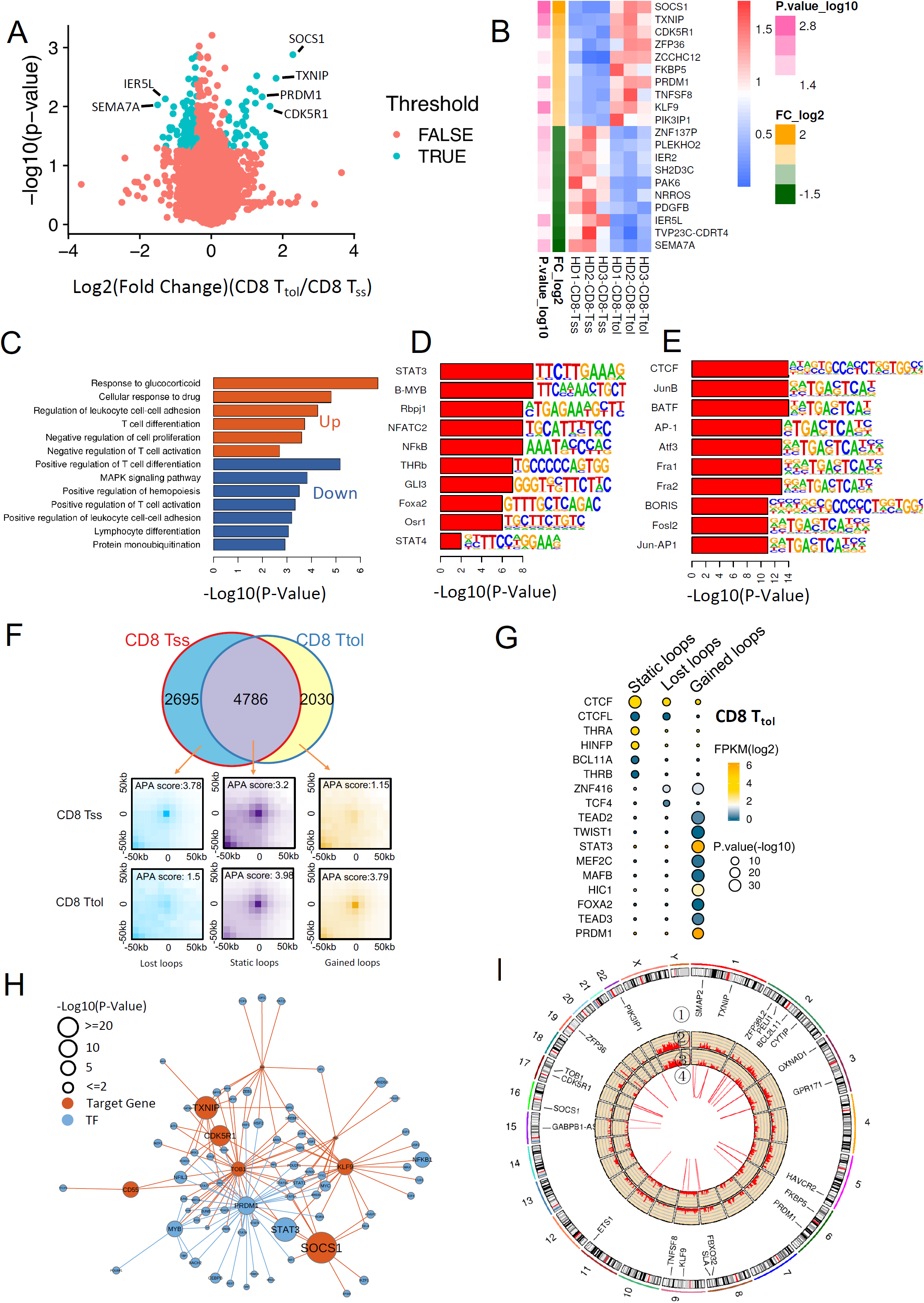
G-CSF regulates target gene expression by chromatin accessibility. **(A)** Volcano plot comparing CD8 T_ss_ and T_tol_. The X-axis is the fold change (log2). Among the genes, 54 genes were significantly upregulated and 78 genes were significantly downregulated. **(B)** Corresponding to A, the top ten genes with high and low expression were identified (red: high expression; blue: low expression). **(C)** Gene pathway enrichment analysis for the upregulated (red) and downregulated (blue) genes in CD8 T_tol_ cells. **(D)** Motif results predicted upregulated chromatin accessibility of CD8 cells by HOMER software. **(E)** Motif results predicted downregulated chromatin accessibility of CD8 cells. **(F)** Venn diagram of CD8 T_ss_ and CD8 T_tol_ chromatin loops. APA analysis was performed on the three types of loops in the Venn diagram to verify the reliability of each type of loop. **(G)** Motif results predicted different types of loops of CD8 T_tol_ cells by HOMER software. CTCF is the most significant transcription factor enriched by static loops. This finding is consistent with the existing literature. **(H)** The regulatory network map of highly expressed genes and enhanced transcription factors in CD8 T_ss_ and CD8 T_tol_. Red dots represent transcription factors, and purple dots represent target genes. The regulatory relationship between transcription factors and genes comes from Yan et al. (Yan et al., 2012). **(I)** Highly expressed genes in the loop anchor region of CD8 T_tol_ compared to CD8 T_ss_. Of the 55 upregulated genes, 29 were located in the anchor region of the loop. From the outer circle to the inner circle: ⍰: gene name; ⍰: gene expression levels from RNA-seq data in CD8 T_tol_; ⍰: chromosome accessibility from ATAC-Seq in CD8 T_tol_; ⍰ red lines: chromatin loops overlap with genes.

We next evaluated the expression of genes and differentially activated TFs of interest. The TF NFAT not only initiates transcriptional programs of T cell activation but also induces T cell tolerance (Kang et al., 1992; Martinez et al., 2015). In CD8^+^ T_tol_, the expression of AP-1 was significantly downregulated, accompanied by the upregulation of NFATC2. In CD4^+^ T_tol_, the expression of Jun was also significantly downregulated, accompanied by the upregulation of NFKBIA (Figure 3A-D and S2B-E). In mouse models, T cell anergy could be induced if the interaction with its transcriptional partner AP-1 (Fos/Jun) was prevented (Kang et al., 1992; Martinez et al., 2015; Sundstedt et al., 1996). Thus, our observations suggest a role for the downregulation of AP-1 (Fos/Jun) in inducing CD4^+^ and CD8^+^ T cell tolerance *in vivo*.

B lymphocyte-induced maturation protein-1 (Blimp-1), a zinc finger–containing transcriptional repressor encoded by PRDM1, could directly inhibit IL-2 transcription and attenuate T cell proliferation both *in vivo* in a mouse model and *ex vivo* (Kallies et al., 2006; Martins et al., 2008; Martins et al., 2006). Consistently, in CD4^+^ T_tol_ and CD8^+^ T_tol_ cells, the expression of PRDM1 was significantly upregulated compared with that of CD4^+^ T_ss_ and CD8^+^ T_ss_ cells, respectively (Figure 3B and Figure S2B). High Blimp-1 has been shown to promote the expression of inhibitory receptors and foster CD8^+^ T cell exhaustion in a mouse model (Shin et al., 2009). Thus, future studies are necessary to understand the role of PRDM1 in inducing human T cell tolerance.

The 3D genome interaction maps of CD4^+^ and CD8^+^ T cells were next investigated (Figure S3C-I) (Li et al., 2018; Wu et al., 2017). The switch of A/B compartments from steady-state T cells to tolerant ones is shown for chromosome 10 in Figure S3G. Approximately 7.4% and 7.3% of the genome regions in CD4^+^ and CD8^+^ T cells, respectively, switched from compartment A to B after tolerance induction. Meanwhile, 6.6% and 4.5% of the genome regions in CD4^+^ and CD8^+^ T cells, respectively, switched from compartment B to A (Figure S3G-I). We found that the length of the TADs decreased in the tolerant CD4^+^ and CD8^+^ T cells compared to the steady-state cells (Figure S4A-D).

We found 7,481 loops in CD8^+^ T_ss_ and 6,186 loops in CD8^+^ T_tol_, with 4,786 loops in both lists (Figure 3F). Compared with those in CD8^+^ T_ss_, STAT3 and PRDM1 motifs are enriched in the gained loop anchors and highly expressed in CD8^+^ T_tol_, which is consistent with the ATAC-seq results (Figure 3D). This suggests that STAT3 may be a structural protein that mediates the gained chromatin loop. The lost loop anchor-enriched TFs include ZNF416 and TCF4 in CD8^+^ T_tol_ (Figure 3G). The number of loops of steady-state and tolerant CD4^+^ T cells was 9787 and 9953, respectively, with 7100 in both lists (Figure S4E). According to the interaction between TFs and target genes (Chen et al., 2018), we plotted the network of upregulated genes and enhanced regulatory transcription factors in CD8 cells (Figure 3H), which shows that SOCS1 is highly expressed and that the strongest TF regulating SOCS1 is STAT3. We also identified a regulatory network between highly expressed genes and enhanced TFs in CD4^+^ cells (Figure 2SE). We demonstrated that more than 53% of upregulated genes, such as SOCS1, PRDM1, and KLF9, are in the loop anchor region of CD8 T_tol_ compared with CD8^+^ T_ss_ (Figure 3I). These observations suggest that chromatin co-accessibility could be determined by the 3D conformation of the genome and may correspond to coordinated regulation of multiple cis- and trans-regulatory elements, including gene expression regulated by known T-cell enhancers and promoters, during *in vivo* induction of human T cell tolerance.

### Association of STAT3 with SOCS1 expression in tolerant T cells

We next explored the regions of spatial interaction with the promoter of SOCS1 and which transcription factors bind to these regions. We showed that the interaction of SOCS1 and the enhancer region in CD8^+^ T_tol_ cells was higher than that in CD8^+^ T_ss_ cells (Figure 4A). In the chromosome 16, the SOCS1 gene is located within one TAD in CD8^+^ T_ss_ (Figure 4B, top). Then, we plotted the chromatin structure, histone modification and TF binding sites around the SOCS1 gene (Figure 4B). The genome-browser view of CTCF and STAT3 binding sites suggests that the CTCF protein mediates the interaction between the SOCS1 locus and the upstream chromatin region, and STAT3 proteins mediate the interaction between SOCS1 and downstream super-enhancers. From Hi-C data, the interaction between the SOCS1 locus and upstream heterochromatin is weakened, and the interaction between SOCS1 and downstream super-enhancers is enhanced in CD8^+^ T_tol_ cells compared to CD8^+^ T_ss_ (Figure 4B) (Wunsche et al., 2018). These results suggest that a new association of STAT3 with SOCS1 expression emerged during the *in vivo* induction of human T cell tolerance. In support of this hypothesis, genome-wide statistics showed that genes with long-range interactions with heterochromatin tended to be expressed at low levels, while genes with long-range interactions with enhancers tended to be highly expressed (Figure 4C).

**Figure 4.**
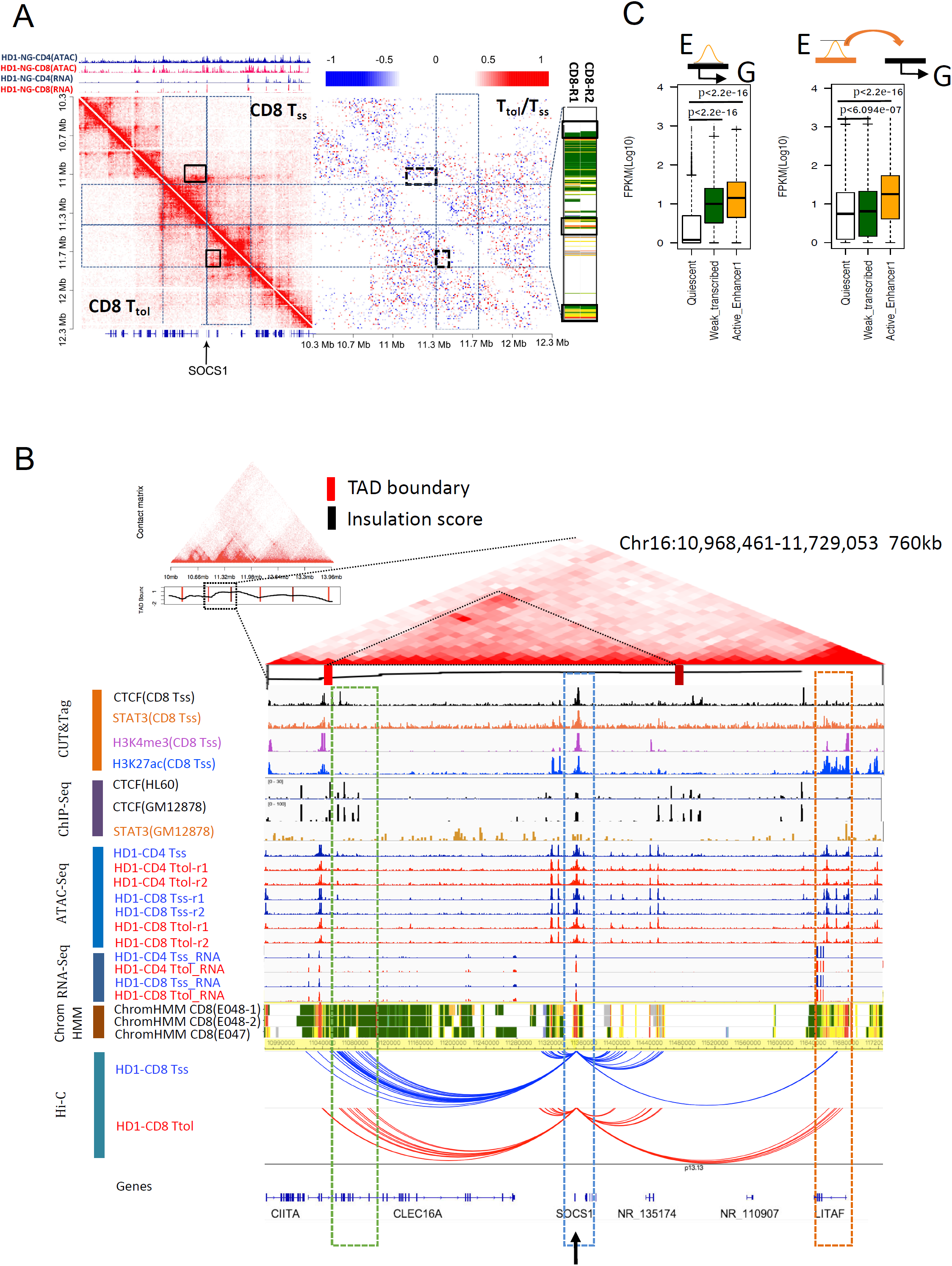
STAT3 regulates SOCS1 expression by 3D genome and chromatin accessibility. **(A)** Similar to 2H, a heatmap showing an example gene with high expression in CD8 T cells located in a differential loop between CD8 T_tol_ cells and CD8 T_ss_ cells on chr16 (chr16: 10,300,000–12,300,000). **(B)** Hi-C interaction matrix of a region (chr16: 10 Mb-14 Mb) in CD8 T_ss_ cells shows the TAD boundaries around the SOCS1 gene. Top: Hi-C interaction matrix, bottom: TAD boundaries (vertical bars) and insulation scores. Genome-browser view of CTCF and STAT3 binding sites, histone modification, chromatin accessibility, gene expression chromatin states and 3D genome interaction around the SOCS1 gene in CD8 T cells. In the chromatin state line, white represents the quiescent chromosome. Green represents chromatin with weak transcription. Yellow represents strong enhancers. The green box represents the region of chromatin with reduced interactions with SOCS1 after G-CSF mobilization. The yellow box represents the region of chromatin with reduced interactions between G-CSF and SOCS1 after G-CSF mobilization. The blue box represents the promoter regions of SOCS1. **(C)** Effects of different chromatin states on gene expression. The boxplot on the left shows the effect of in situ chromatin status on genes. The boxplot on the right represents the effect of chromatin interactions on long-distance gene expression. Enhancers activate gene expression more than the other two chromatin states.

To investigate whether there is colocalization between STAT3 and CTCF (Vahedi et al., 2012; Weintraub et al., 2017), we performed ChIP-seq and CUT&Tag experiments. For example, many of the binding sites of STAT3 and CTCF are colocalized in and around SOCS1 (Figure 5A) and TXNIP (Figure 5B) which are upregulated after G-CSF mobilization. Further, STAT3 and CTCF colocalized in the whole genome of human CD8 T cells and GM12878 cell lines (Figure 5C-F and Figure S5A-D). There was a significant overlap between the CTCF peaks and the STAT3 peaks in CD8 cells, as shown by the Venn diagram (*P*<1e-10, Figure 5F). The peaks of STAT3 binding are classified into promoter region and the enhancer region (Figure 5G-5H), and there is a significant spatial interaction between the promoter region and enhancers (Figure 5I-5J). These results strongly suggest that the STAT3 complex is involved in the enhancer and promoter interaction (Figure 5G-5J and Figure S5E-G). Consistent with our observation, STAT4 binding in the genome contributes to the specification of the nuclear architecture around *Ifng* during Th1 differentiation (Hakim et al., 2013). Furthermore, we observed both CTCF and STAT3 foci in the nuclei of Jurkat cells by immunofluorescence staining (Figure S5H). Collectively, these results suggest that a STAT3-mediated enhancer-promoter interaction induces SOCS1 expression during the *in vivo* induction of T cell tolerance (Figure 5K). This example highlights a potentially important way in which T cell function could be modulated.

**Figure 5.**
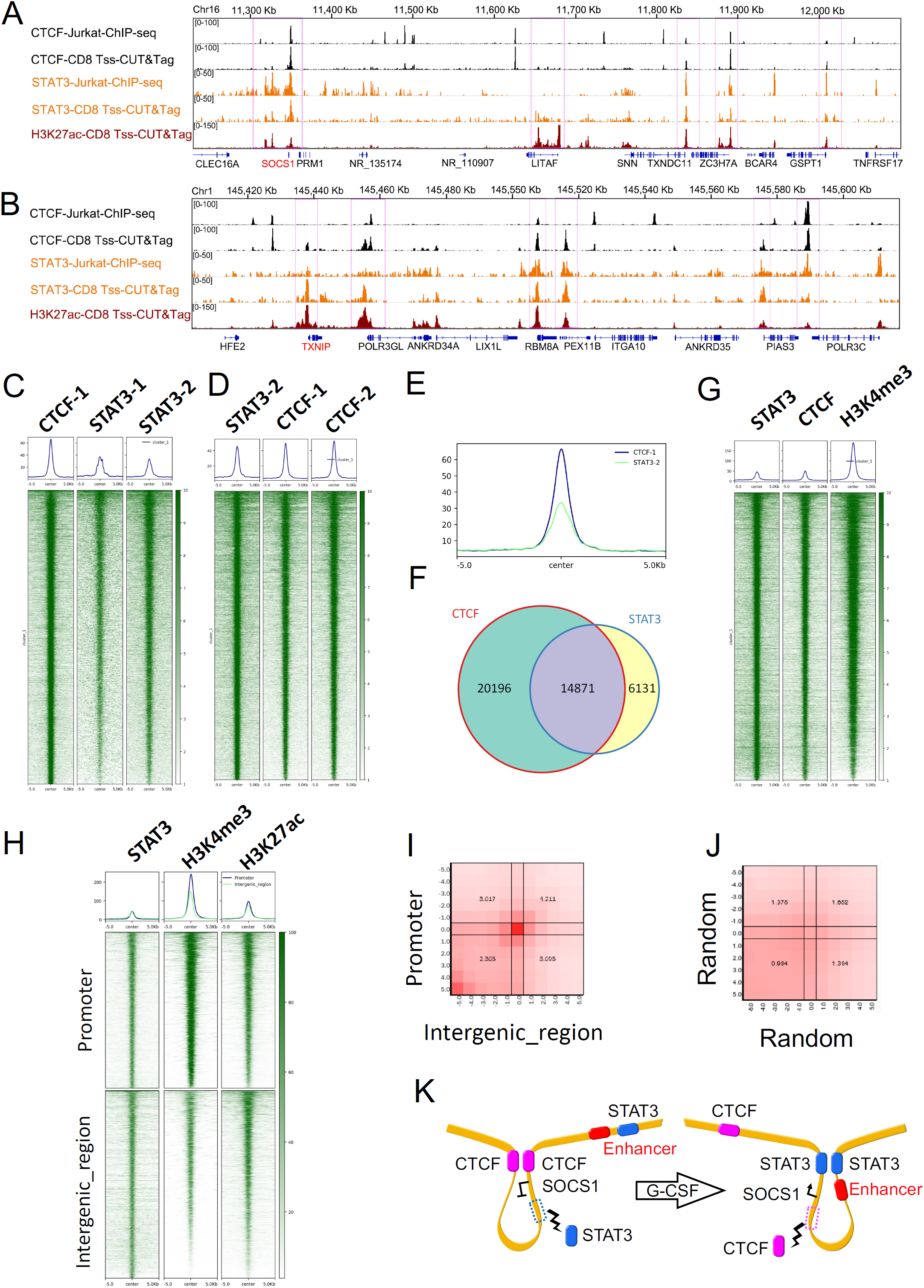
STAT3 and CTCT are colocalized in the whole genome, and STAT3 mediates the spatial interaction between enhancers and promoters. **(A-B)** The UCSC browser views showing histone modifications and transcription factor (TF) binding sites of SOCS1(A) and TXNIP (B) in Jurkat and CD8 T_ss_ cells using ChIP-seq and CUT&Tag data. Pink dotted box represents where two transcription factors are co-localized **(C)** Heatmaps displaying STAT3 and CTCF colocalized in the whole genome in CD8 T_ss_ cells using CUT&Tag data (the top 5000 CTCF peaks in CD8 T_ss_ cells). **(D)** Heatmaps displaying STAT3 and CTCF colocalized in the whole genome in CD8 Tss cells using CUT&Tag data (the top 5000 STAT3 peaks in CD8 T_ss_ cells). **(E)** Aggregate plot of CTCF binding (blue line) and STAT3 binding (green line) at ± 5.0 Kb from the CTCF peaks in CD8 cells. **(F)** Venn diagram showing the overlap between CTCF peaks (red) and STAT3 peaks (blue) in CD8 T_ss_ cells. p < 1e−10, hypergeometric test. **(G)** Heatmaps displaying STAT3 occupancy and active promoters (the top 5000 STAT3 peaks in CD8 T_ss_ cells). **(H)** The peaks of STAT3 binding are classified into two clusters. The first cluster is the promoter region, which overlaps with the promoter of all genes (± 1 kb around the transcription start site), and the second is the intergenic region. **(I)** Heatmap of the interaction between the promoter region and the intergenic region in the spatial interaction between enhancers and promoters. **(J)** A random selection of the same number of enhancers and promoter peaks has no significant spatial interaction. **(K)** The model of SOCS1 regulation during the *in vivo* induction of human T cell tolerance.

### *In vitro* experiments identify the role of SCOS1 in modulating T cell function

The upregulation of SOCS1 during the *in vivo* induction of human T cell tolerance (Figures 3B and S4B; Figure 4A-C) (Jun et al., 2004; MacDonald et al., 2014; Pan et al., 1995) prompted us to investigate the effects of SOCS1 overexpression on T cell function *in vitro*. We overexpressed SOCS1 in steady-state T cells by lentivirus and found that the SOCS1 expression level was increased approximately 30-fold in the SOCS1 OE group (Figure 6A). High expression of SOCS1 inhibited T cell proliferation, and most T cells in the SOCS1 OE group were blocked in the G0 stage compared to those in the CT or non-infection group (Figure 6B). The proliferative ability of CD4^+^ T cells was decreased in the SOCS1 OE group compared to the CT or non-infection group, while CD8^+^ T cells had no change (Figure 6C-D, Figure S6A). Moreover, highly expressed SOCS1 in T cells also promoted TIGIT expression (Figure 6E, Figure S6B-C). There were no significant differences in the secretion of cytokines, such as IFN-γ, IL-2, IL-17, IL-4, and IL-10, of CD4^+^ T and CD8^+^ T cells between the SOCS1 OE group and CT group (Figure 6F, Figure S7A-C).

**Figure 6.**
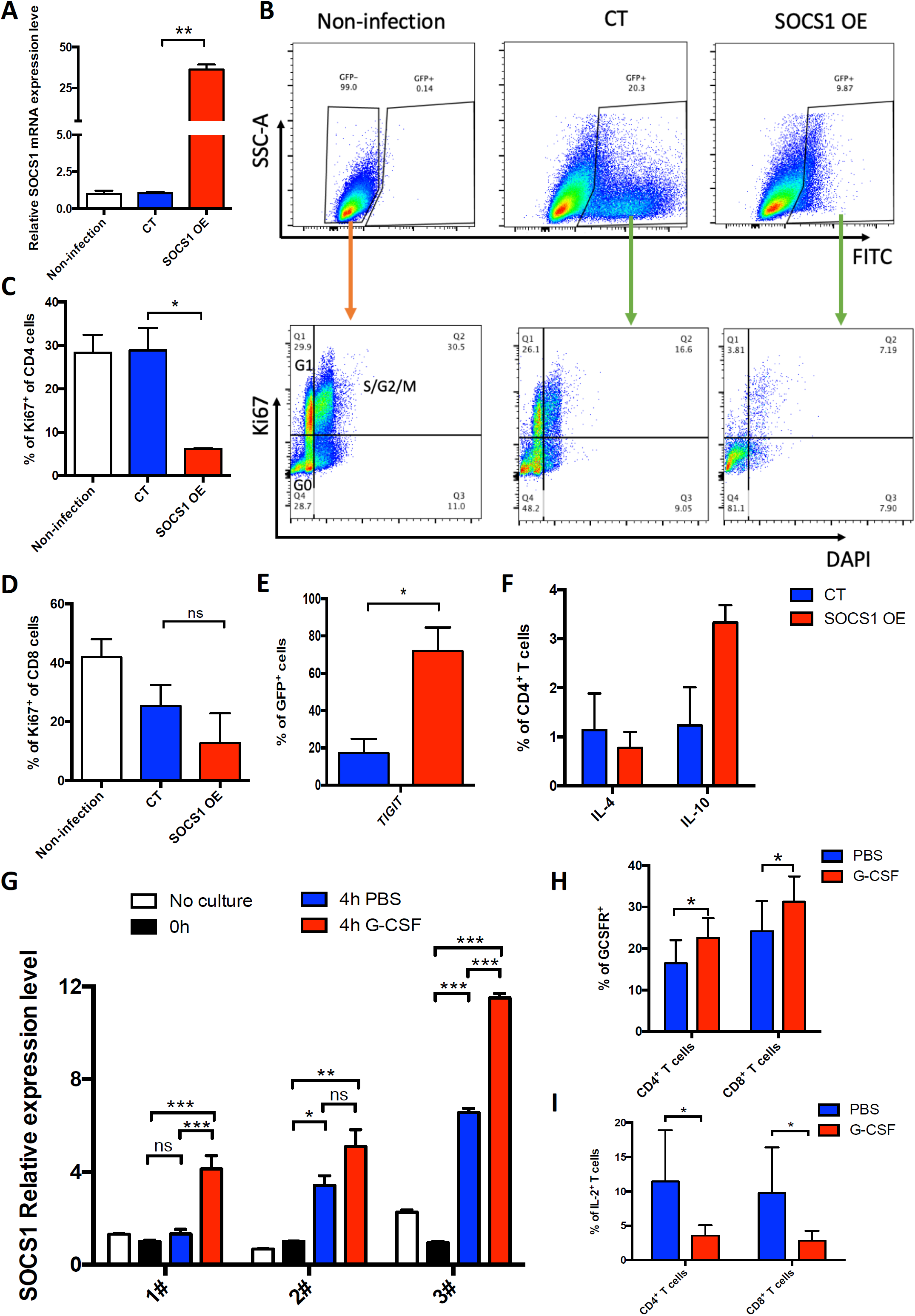
High expression levels of SOCS1 impaired T cell proliferation *in vitro*. **(A)** SOCS1 was overexpressed by lentivirus in CD3^+^ T cells from healthy donor bone marrow. Quantitative real-time RT-PCR was used to detect SOCS1 expression levels. Error bars represent the mean ± SEM values from 3 independent experiments, **p<0.01. **(B)** Flow cytometric analysis of proliferation in GFP^+^ cells. **(C-D)** Percentage of Ki67^+^ cells in CD4^+^ cells (C) or CD8^+^ cells (D). Error bars represent the mean ± SEM values from 3 independent experiments, *p<0.05. **(E)** The expression level of TIGIT detected by flow cytometry. Error bars represent the mean ± SEM values, *p<0.05. **(F)** IL-4 and IL-10 secretion levels in CD4^+^ T cells of GFP^+^ cells. Error bars represent the mean ± SEM values. **(G)** The expression level of SOCS1 in G-CSF-stimulated CD3^+^ T cells at 4 h from 3 independent healthy donors. Error bars represent the mean ± SEM values from 3 independent experiments. One-way ANOVA, *p<0.05, **p < 0.01, ***p < 0.001. **(H)** Representative GCSFR (CD114) expression level of CD3^+^ T cells upon G-CSF treatment at 72 h *in vitro*. Data are expressed from 3 independent healthy donors. Error bars represent the mean ± SEM values from 3 independent experiments, *p<0.05. **(I)** IL-2 secretion in G-CSF-stimulated CD3^+^ T cells at 72 h from 7 independent healthy donors.

We next overexpressed SOCS1 in the Jurkat T cell line and performed Western blotting to detect the JAK-STAT signaling pathway. The results showed that high SOCS1 expression inhibited the phosphorylation of STAT3 (Figure S7D). This finding partially explains why SOCS1 overexpression inhibits T cell proliferation mainly by inhibiting the JAK-STAT signaling pathway. These data suggested that a high expression level of SOCS1 could promote T cell behavior as a tolerance-like phenotype *in vitro*. To further prove this hypothesis, we purified CD3^+^ T cells from G-CSF-treated healthy donors and knocked down SOCS1 expression by siRNA. Of the three siRNA sequences tested, two (siRNA-1, siRNA-2) were found to be effective at reducing SOCS1 mRNA expression as assessed by qRT-PCR (Figure S8A). Then, we investigated the effect of SOCS1 silencing on cytokine production in T cells. Decreased SOCS1 expression resulted in lower levels of IL-10 secretion, as evidenced by FACS and ELISA, than those in the NC control group, but there was no statistically significant difference (Figure S8B-D). The above results could be explained by the fact that SOCS1 expression could not be completely knocked down by siRNA *in vitro*; therefore, *in vivo* experiments with SOCS1-specific knockout in T cell mouse models are needed.

Considering that this study is based on the platform of T cell tolerance induced by G-CSF, the direct relationship between G-CSF and SOCS1 was assessed with highly purified CD3^+^ T cells from 7 independent heathy donor BM samples *in vitro*. We found that G-CSF stimulation *ex vivo* led to a peak of SOCS1 mRNA production after 4 h of culture, followed by recovery after 72 h of culture (Figure 6G, Figure S9A-B). After 72 h of culture of CD3^+^ T cells with G-CSF stimulation *in vitro*, the G-CSFR expression level was significantly increased (Figure 6H, Figure S9C). Moreover, IL-2 was decreased in the G-CSF treatment group after 72 h of culture, which may indicate that G-CSF suppressed the differentiation of T cells to the Th1 type *in vitro* (Figure 6I, Figure S9D). Other cytokines, such as IFN-γ, IL-17, IL-4, and IL-10, were not observed to change (Figure S9E-H).

## DISCUSSION

Here, we report a method charting the epigenomic and transcriptomic landscape of human BM CD4^+^ and CD8^+^ T cells during the *in vivo* induction of immune tolerance (Jun et al., 2004; Rutella et al., 2005). Unsupervised clustering of accessible chromatin regions, specifically distal elements, groups individual cell types with high clustering purity, suggesting that these distal regulatory elements precisely define T cell immunological characteristics during induction of tolerance. In addition, changes in the 3D genome compartment status might influence the accessibility of genomic regions to transcription factors or other regulatory proteins, which could be particularly important for certain subsets of genes. Integration of multiomic data enabled us to identify a novel regulatory model for SOCS1 expression during T cell tolerance induction. Furthermore, we identified the role of SOCS1 in inducing T cell tolerance *in vitro*. The methodologies developed here might have important implications for addressing immunological profiles, such as those of dendritic cells and B cells, in other contexts of tolerance induction (Lynn et al., 2019; Nemazee, 2017) as well as in cellular therapy.

In mouse models, deletion of NR4A1, Blimp-1, Cbl-b, NFAT, TSC1, GRAIL, or Egr-2 impairs induction of T cell tolerance *in vivo* (Haymaker et al., 2017; Jeon et al., 2004; Liu et al., 2019; Macian et al., 2002; Martins et al., 2008; Xie et al., 2012; Zheng et al., 2012). Overexpression of c-Jun in T cells renders them resistant to exhaustion (Lynn et al., 2019). In tolerant human CD4^+^ and CD8^+^ T cells, we observed increases in NFAT1, Blimp-1, and NFKBIA as well as a decrease in Jun, AP-1, and Fos (Soto-Nieves et al., 2009), suggesting an association of these genes and TFs with immune tolerance. However, there were significant differences in the expression of some genes and TFs, such as NR4A1 and Egr-2, between human and mouse tolerant T cells. Several reasons account for the difference. First, the species of T cells are different. Second, different methods, for example, blockade of costimulatory signals in mice by others (Liu et al., 2019) and treatment of healthy donors with G-CSF in our study (Jun et al., 2004), for tolerance induction via different signaling pathways might lead to differences in gene and TF expression (Liu et al., 2019; Soskic et al., 2019; Zhu and Paul, 2010). Overall, our study not only confirmed tolerance-inducing genes and TFs in human tolerant T cells, which were first observed in a mouse model, but also provided a platform to find novel genes, such as KLF-9.

In this study, we found that both tolerant CD4^+^ T cells and CD8^+^ T cells are distinguished by high expression of SOCS1 (Davey et al., 2005). Based on the landscape of the 3D genome structure and chromatin accessibility in CD4^+^ and CD8^+^ T_tol_ cells, we observed a STAT3-mediated enhancer/promoter interaction for SOCS1 gene expression and proposed a novel model in which STAT3 could replace CTCF and form new chromatin loops, leading to the expression of SOCS1 (Figure 5I). Our results are consistent with a study showing that NF-κB could compete with CTCF, forming a new loop and enhancing PD-L1 expression (Chen et al., 2018). Several studies have shown that SOCS1 negatively regulates the activation of the STAT/Jak signaling pathway (Davey et al., 2005; Yang et al., 2013), suggesting that it could be a potent gene in inducing tolerance. In this study, *in vitro* experiments using human primary T cells showed that overexpression of SOCS1 led to a significant suppression of T cell proliferation and upregulation of the inhibitor immune receptor, suggesting that SOCS1 can lead to T cell tolerance, although *in vivo* studies are needed to confirm the role of SOCS1 in inducing tolerance and the underlying mechanisms.

In summary, based on *in vivo* induction of a human T cell tolerance model (Jun et al., 2004; Rutella et al., 2005) and multiomic analyses, we established a platform for discovering novel genes and TFs that induce immune tolerance. Our data resource will serve as a valuable tool for the community to further elucidate the gene regulatory networks controlling the induction of human T cell tolerance. In addition, the data on SOCS1 pathways in orchestrating the characteristics of tolerogenic T cells, together with recent findings on the role of SOCS1 in intestinal immune tolerance, suggest that this gene could be a suitable target for tolerance induction.

## ACKNOWLEDGEMENTS

This work was partly supported by grants from the National Key Research and Development Program of China (2017YFA0104500), the Beijing Municipal Science and Technology Commission (Z181110009618032), the National Natural Science Foundation of China (Grant Nos. 81670168, 81621001, 81530046, and 81770189), and the Key Program of the National Natural Science Foundation of China (Grant No. 81230013). The CAMS Innovation Fund for Medical Sciences (CIFMS) (Grant No. 2019-I2M-5-034) and the Fundamental Research Funds for the Central Universities also provided funding. R.L and C.L were supported by the National Natural Science Foundation of China (31871266), the Chinese National Key Research and Development Program of China (2016YFA0100103), and NSFC Key Research Grant 71532001. Part of the data analysis was performed on the High Performance Computing Platform of the Center for Life Sciences, Peking University. We thank Shanghai Jiayin Biotechnology, Ltd., for their help and suggestions in the ChIP-seq and CUT&Tag experiments. We thank all the faculty members who participated in these studies.

## AUTHOR CONTRIBUTIONS

Contribution: X.-J.H., C.L., and Y.-J.C. designed the study; R.-F.L. and H.-D.G. collected data; R.-F.L., H.-D.G., Y.-J.C., C.L., and X.-J.H. analyzed the data and drafted the manuscript; all authors contributed to data interpretation and manuscript preparation and approved the final version.

## CONFLICT OF INTEREST

The authors declare no conflicts of interest.

## STAR METHODS

### T cell isolation and culture

Human bone marrow mononuclear cells (BMMCs) were isolated from the BM of healthy donors before and after *in vivo* G-CSF application by Ficoll density centrifugation. CD3^+^ T cells were purified by positive selection (CD3 MACS MultiSort beads; Miltenyi Biotec, Bergische Gladbach, Germany). The isolated CD3^+^ T cells were cultured in IMDM medium (Gibco, Invitrogen, Carlsbad, CA) containing 10% BIT 9500 (Stemcell Technologies, Vancouver, CA) and stimulated with Dynabeads Human T-Activator CD3/CD28 (Gibco, Invitrogen, Carlsbad, CA). The study was approved by the Institutional Review Board of Peking University. Written informed consent was obtained from all healthy donors in accordance with the Declaration of Helsinki.

### Hi-C experiments

The cells were resuspended in fresh PBS. Cell counts were performed. Then, a cell suspension with a final concentration of 1×10^6^ cells per 1 ml of PBS was prepared. A total of 1 × 10^6^ cells were isolated and crosslinked with 1% formaldehyde for 10 min at room temperature, and then, 2.5 M glycine solution was added to a final concentration of 0.2 M. Then, the cells were collected, flash-frozen in liquid nitrogen and stored at −80 °C. The Hi-C experiment was performed following the in situ Hi-C protocol (Rao et al., 2014).

### RNA-seq experiments and analysis

Total mRNA with a polyA tail was extracted and reverse transcribed to cDNA for sequencing. Three biological repeats were performed for each sample, and 20 million reads were sequenced for each repeat. The sequenced reads were mapped to the human reference genome (hg19) by TopHat2 (Kim et al., 2013), and gene expression was quantified by Cufflinks (Trapnell et al., 2010). We used RStudio software for the downstream statistical analyses.

### ATAC-seq experiments and analysis

The ATAC-seq experiment was performed following Buenrostro’s protocol (Buenrostro et al., 2013). Two biological repeats were used for each sample, and 20 million reads were sequenced for each repeat. The sequenced reads were mapped to the human reference genome (hg19) by Bowtie2 (Kim et al., 2013), and peak signals were quantified by MACS2 and deepTools. We used RStudio software for the downstream statistical analyses.

### Hi-C data analysis

We performed read mapping and filtering of the Hi-C data following previous methods (Jin et al., 2013). All Hi-C sequencing reads were mapped to the human reference genome (hg19) using Bowtie2 (Langmead and Salzberg, 2012). The two ends of paired-end reads were mapped independently using the first 36 bases of each read. We filtered out redundant and nonuniquely mapped reads and kept the reads within 500 bp upstream of enzyme cutting sites (Mbol) due to size selection. We used the iterative correction and eigenvector decomposition (ICE) method (Imakaev et al., 2012) to normalize raw interaction matrices.

### A/B compartment analysis

We used ICE-normalized interaction matrices at 500 kb resolution to detect chromatin compartment types by the R package HiTC (Servant et al., 2012). Positive or negative values of the first principal component separated the chromatin regions into two spatially segregated compartments. The compartment with a higher gene density was assigned the A compartment, and the other compartment was assigned the B compartment (Barutcu et al., 2015).

### TAD analysis

We used ICE-normalized interaction matrices at 40 kb resolution to call TAD by a Perl script matrix2insulation.pl (https://github.com/blajoie/crane-nature-2015). A higher resolution was used because TADs are smaller than A/B compartments. Insulation scores (IS) were calculated for each chromosome bin, and the valleys of the IS identified the TAD boundaries. TADs smaller than 200 kb or located in telomeres/centromeres were filtered out as in previous methods (Crane et al., 2015). In comparisons of TADs between two cell lines, at least 70% overlap between two TADs was considered conserved TADs (Taberlay et al., 2016). We used BEDtools with the option of “intersectBed −f 0.70 – r” to identify conserved TADs (Quinlan and Hall, 2010).

### Gene ontology analysis

We used DAVID Bioinformatics Resources 6.7 for gene ontology analysis (Huang et al., 2009). All human genes were used as the background gene list.

### Chromatin immunoprecipitation

Jurkat cells were fixed in 1% formaldehyde (Sigma-Aldrich, F8775) for 10□min at 37□°C. Subsequently, glycine was added to 125□mM and incubated at 37□°C for 5□min at 37□°C. Next, the cells were pelleted and washed twice with cold PBS. The pellets were stored at −80□°C until use.

Nuclei from 10□M cells per ChIP-seq were extracted, and chromatin was sonicated with a Bioruptor Sonication Device. Immunoprecipitation reactions were performed overnight with STAT3 (Cell Signaling Technology, 9139S, MA), H3K27ac or CTCF (ABclonal, A1133, China) antibodies. The next morning, antibodies and chromatin were captured using Protein G Dynabeads (Thermo Fisher). The material was washed, eluted and treated with RNase A for 30□min at 37□°C and proteinase K for 3□h at 65□°C.

### Library preparation and sequencing

Library preparation of ChIP-seq DNA was performed using the Ultra II Library Prep Kit (NEB E7103L) and Multiplex Oligos for Illumina (NEB E7335L) and sequenced on an Illumina NextSeq 2500 (150□base pairs single end).

### ChIP–seq data processing, heat map generation, and edgeR analysis

H3K27ac, CTCF, and STAT3 ChIP–seq analyses were performed with an average range of 20–25 × 10^6^ reads per independent ChIP–seq experiment. ChIP–seq reads were mapped to the hg19 genome with Bowtie2 using default parameters. Aligned reads were filtered for a minimum MAPQ of 30, and duplicates were removed using SAMtools. Signal tracks were generated by first using BEDTools to produce bedGraph files scaled to 10 million reads per data set. Then, the UCSC Genome Browser utility bedGraphToBigWig was used with default parameters to generate bigwig files. Peaks were called using MACS2 with default parameters. Heat maps of ChIP–seq signal profiles were generated with the HOMER (http://biowhat.ucsd.edu/homer/index.html) tool annotatePeaks with the following parameters: -ghist 50, -size 10000. ChIP–seq peaks exhibiting differential H3K27ac or STAT3 signals across the time course were identified using edgeR similar to the process described above.

### CUT&Tag experiments and analysis

The CUT&Tag experiments were performed as previously described (Kaya-Okur et al., 2019) (Vazyme TD901 kit) to generate DNA libraries derived from human CD8 Tss cells. We use the SEACR peak caller (http://seacr.fredhutch.org), which was expressly designed for CUT&RUN and CUT&Tag data, to call peaks.

### *In vitro* stimulation with G-CSF

Isolated CD3^+^ T cells from healthy donors were incubated with G-CSF (100 ng/ml) for 4 h or 72 h at 37 °C and 5% CO_2_.

### Lentivirus-mediated SOCS1 overexpression in T cells

The SOCS1-overexpressing lentivirus was purchased from Sangon Biotech (Shanghai, China). CD3^+^ T cells were prestimulated for 24 h with Dynabeads Human T-Activator CD3/CD28 in IMDM medium containing 10% BIT 9500, and rhIL-2 was added at a dose of 100 U/ml. After 24 h, the cells were transduced with thawed lentiviruses that were added directly to the plate. Then, 6 μg/ml polybrene (Sigma, USA) was added. The cells were incubated for another 24 h at 37 °C and 5% CO_2,_ and fresh medium was changed. GFP^+^ cells were isolated after a 72-h infection and cultured in IMDM medium containing 10% BIT 9500 with rhIL-2 routinely used.

### T cell transfection by siRNA oligo

Bone marrow samples were obtained from 6 healthy donors after treatment with recombinant G-CSF at a dosage of 5 μg/kg/d for 5-6 consecutive days. CD3^+^ T cells were isolated from the bone marrow of post-G healthy donors, cultured in IMDM containing 10% BIT 9500 with rhIL-2 100 U/ml, and stimulated with Dynabeads Human T-Activator CD3/CD28. After 24 h of culture, the CD3^+^ T cells were transfected with 21 base-pair siRNA oligonucleotides (siRNA-1, 5’-CCAGAACCTTCCTCCTCTT-3’; siRNA-2, 5’-ACACGCACTTCCGCACATT-3’; siRNA-3, 5’-CTGGGATGCCGTGTTATTT-3’). Then, 2.5 μl of 20 μM oligonucleotides was added to 5 μl of Lipo-3000 (Invitrogen, Carlsbad, CA) and 42.5 μl of serum-free OPTI-MEM (Gibco, Invitrogen, Carlsbad, CA) and incubated at 25 °C for 20 min. Then, 50 μl of the mixture was added to each well of CD3^+^ T cells and incubated for 48 h at 37 °C. After 48 h of culture, the cells were collected, and the knockdown efficiency was detected by RT-PCR.

### Flow cytometric analysis

Surface staining was performed with directly conjugated monoclonal antibodies for 20 min at room temperature. The cells were washed and resuspended in phosphate-buffered saline (PBS) before flow cytometric analysis. The monoclonal antibodies used were anti-human CD4-Percp-Cy5.5/APC-H7, CD8-APC-R700/V500, PD-1-PE-Cy7, Tim-3-APC, Tigit-BV605, 2B4-AF700, and CD160-PE (BD Bioscience San Diego, CA, USA). Intracellular staining was carried out by using a fixation/permeabilization kit (BD Bioscience) after resuspension according to the manufacturer’s instructions. Ki-67-PE (BD Pharmingen) was added and incubated for 20 min at room temperature.

### Cytokine detection by flow cytometry

T cells were stimulated with Dynabeads Human T-Activator CD3/CD28. After 72 h of culture, Golgiplug (BD Pharmingen, San Diego, CA, USA) was added for 4 h. Cells were harvested for surface staining as described above. Intracellular staining was carried out by using a fixation/permeabilization kit (BD Bioscience) after resuspension according to the manufacturer’s instructions. IL-2-V450, IFN-γ-BV510, IL-17-PE, IL-4-APC, and IL-10-PE (BD Pharmingen) were added and incubated for 20 min at room temperature.

### RT-PCR

RNA was extracted using the RNeasy Mini Kit (Qiagen, 74106) according to the manufacturer’s protocol. For quantitative PCR, first strand synthesis was performed using a cDNA reverse transcription kit (TaKaRa, RR047A) according to the manufacturer’s protocol. Quantitative PCR assays were performed in 96-well MicroAmp Fast Optical 96-Well Reaction Plates (Applied Biosystems, 4344904) using SYBR Green (Roche, 04913914001). Signals were detected using a 7500 Real-Time PCR System (Applied Biosystems). Target gene cycle numbers were normalized to the housekeeping gene 18S to obtain a ΔCT value. The 2^-ΔΔCT^ method was used. The primer sequences were as follows: SOCS1 forward: 5’-CACGCACTTCCGCACATTC-3’; SOCS1 reverse: 5’-TAAGGGCGAAAAAGCAGTTCC-3’; human 18S forward: 5’-ACCGATTGGATGGTTTAGTGAG-3’; and human 18S reverse: 5’-CCTACGGAAACCTTGTTACGAC-3’.

### Code and data

All essential codes used for the analysis are available at GitHub (https://github.com/ ChengLiLab/T_cell_tolerance). The raw sequencing data generated by this project were deposited at Genome Sequence Archive (GSA, http://gsa.big.ac.cn) with accession number PRJCA002316.

## SUPPLEMENTAL FIGURE LEGENDS

**Figure S1. Quality control for sequencing data**

**(A)** Summary of the sequencing data of Hi-C. **(B)** Genome browser view of the ATAC-seq signal around the MDM4 promoter in CD4 and CD8 cells. This image shows that our repeated experiments are highly consistent. **(C)** Insert size histogram for all reads in the HD1 CD8 T cells. **(D)** Unsupervised hierarchical clustering of the ATAC-seq signal of the accessible chromatin regions in the genome. **(E)** Distribution of the accessible chromatin regions in the genome of the HD1 CD8 T_ss_ cells. **(F)** Read count per million mapped reads around the transcription start site (TSS). **(G)** Visualization of the read coverage at transcription start sites by a heatmap.

**Figure S2. Differentially expressed genes and differentially activated TFs in the CD4**^**+**^ **cells before and after mobilization.**

**(A)** Volcano plot comparing CD4 T_ss_ and CD4 T_tol_. The X-axis is the fold change (log2). Among the genes, 52 genes were significantly upregulated and 85 genes were significantly downregulated. **(B)** Corresponding to map A, the top ten genes with high and low expression were identified (red: high expression; blue: low expression). **(C)** Motif results predicted upregulated chromatin accessibility of the CD4 cells using ATAC-seq data by HOMER software. **(D)** Motif results predicted downregulated chromatin accessibility of the CD4 cells using ATAC-seq data by HOMER software. **(E)** The regulatory network map of the highly expressed genes and enhanced transcription factors in CD4 before and after mobilization. Red dots represent transcription factors, and purple dots represent target genes. The regulatory relationship between transcription factors and genes comes from Yan et al. (Yan et al., 2012).

**Figure S3. Specific gene and chromatin structures in the CD4 and CD8 cells after mobilization.**

**(A)** Volcano plot comparing CD4 T_tol_ and CD8 T_tol_ cells. The X-axis is the fold change (log2) of CD4 T_tol_ /CD8 T_tol_. There were 115 genes overexpressed in CD4 T_tol_ cells and 214 genes overexpressed in CD8 T_tol_ cells. **(B)** Corresponding to A, the top ten genes with high and low expression were identified (red: high expression; blue: low expression). **(C)** Whole genome Hi-C interaction matrix of the HD1 CD4 T_ss_ cells. **(D)** Hi-C interaction matrix of the HD1 CD4 T cells on chromosome 16. **(E)** Scatter plots of Hi-C detected the genome interactions of CD8 T cells between two healthy donors. **(F)** Scatter plots of Hi-C detected the genome interactions of CD8 cells before and after G-CSF mobilization. **(G)** Genome-wide proportions of A/B compartment changes among two cells before and after G-CSF mobilization (Fisher’s exact test < 2.2e-16). **(H)** A/B compartments of different cell types. The A/B compartments of chromosome 10 inferred from Hi-C data of various samples. **(I)** Boxplots of expression changes of genes grouped by their A/B compartment changes (t-test).

**Figure S4. TAD and loop structures are influenced by G-CSF.**

**(A)** Examples of conserved and changed TADs (indicated by the red points) in a region (chr16: 1 Mb-19 Mb). **(B)** The length of the TADs in CD4 and CD8 cells (t-test). **(C)** Read count per million mapped reads around the TAD boundaries using CTCF, H3K27ac ChIP-seq and ATAC-seq data. **(D)** Hi-C interaction matrix of the CD8 cells of part of chromosome 16. Black dotted lines represent TADs. **(E)** Venn diagram of the CD4 T_ss_ cells and the CD4 T_tol_ cells in chromatin loops. APA analysis was performed on the three types of loops in the Venn diagram to verify the reliability of each type of loop.

**Figure S5. STAT3 and CTCT are colocalized in the whole genome in the GM12878 and Jurkat cell lines.**

**(A)** Heatmaps displaying STAT3 and CTCF colocalized in the whole genome (the top 5000 CTCF peaks) in GM12878. **(B-C)** Heatmaps displaying STAT3 occupancy at enhancers (B) and active promoters (C) (the top 5000 STAT3 peaks) in GM12878. **(D)** The peaks of STAT3 binding are classified into three clusters. The first cluster includes both enhancer and promoter signals. The second kind of enhancer signal is stronger. The third kind of promoter signal is relatively strong. **(E)** Signal strength of the three kinds of peaks. **(F)** Heatmap of the interaction between Cluster 2 and Cluster 3 in space. The spatial interaction between enhancers and promoters. **(G)** A random selection of the same number of enhancers and promoter peaks has no spatial interaction. **(H)** Immunofluorescence staining of CTCF and STAT3 in Jurkat cells.

**Figure S6. High expression levels of SOCS1 promote exhaustion marker expression.**

**(A)** Strategy of FACS analysis in the SOCS1-overexpressing T cells. **(B)** Flow cytometric analysis of TIGIT and Tim3 expression levels in the SOCS1-overexpressing CD3^+^ T cells. **(C)** Statistical results of the exhaustion marker 2B4, CD160, PD-1, TIGIT, and Tim-3 expression levels in the SOCS1-overexpressing CD3+ T cells. Error bars represent the mean ± SEM values from 3 independent experiments from 3 healthy donors, *p<0.05.

**Figure S7. Cytokine secretion level in T cells after SOCS1 overexpression *in vitro*.**

**(A-B)** Flow cytometric analysis (A) and statistical results (B) of the IL-2, IFN-γ, and IL-17 secretion levels in the SOCS1-overexpressing CD4^+^ T cells or CD8^+^ T cells. Error bars represent the mean ± SEM values from 3 independent experiments from 3 healthy donors. **(C)** Flow cytometric analysis of the IL-4 and IL-10 secretion levels in the SOCS1-overexpressing CD4^+^ T cells. Error bars represent the mean ± SEM values from 3 independent experiments from 3 healthy donors. **(D)** Western blot analysis of the STAT3 and phosphorylated STAT3 levels in the SOCS1-overexpressing Jurkat cell line.

**Figure S8. IL-10 secretion in tolerant T cells *in vitro* after decreasing SOCS1 expression.**

**(A)** SOCS1 knockdown by siRNA in CD3^+^ T cells from post-G healthy donors. The relative expression level of SOCS1 was detected by quantitative real-time RT-PCR. Error bars represent the mean ± SEM values from 6 independent post-G healthy donors, ***p < 0.001. **(B-C)** The IL-10 secretion level was detected by flow cytometry in CD4^+^ T cells (B) or CD8^+^ T cells (C) after SOCS1 was knocked down. Error bars represent the mean ± SEM values from 6 independent post-G healthy donors. **(D)** The IL-10 secretion level was detected by ELISAs in CD3^+^ T cells after SOCS1 was knocked down. Error bars represent the mean ± SEM values from 6 independent post-G healthy donors.

**Figure S9. G-CSF elevated the SOCS1 expression levels and suppressed the Th1 phenotype in T cells *in vitro*.**

**(A)** The SOCS1 expression level was determined by quantitative real-time RT-PCR upon G-CSF treatment *in vitro* at the indicated time points: 4 h, 8 h, 24 h, and 72 h. **(B)** SOCS1 expression levels were determined by quantitative real-time RT-PCR upon G-CSF treatment *in vitro* for 72 h. The SOCS1 expression level was normalized to the PBS control. Error bars represent the mean ± SEM values from 3 independent healthy donors, *p<0.05. **(C)** A representative GCSFR (CD114) expression level of CD3^+^ T cells was determined by flow cytometry upon G-CSF treatment at 72 h *in vitro*. **(D)** Flow cytometric analysis of IL-2 secretion in G-CSF-stimulated CD3^+^ T cells at 72 h from 7 independent healthy donors. (E-F) IFN-γ and IL-17 secretion levels of the CD4+ T cells **(E)** or the CD8^+^ T cells **(F)** after G-CSF stimulation for 72 h *in vitro*. Error bars represent the mean ± SEM values from 3 independent healthy donors. **(G-H)** IL-4 and IL-10 secretion levels of the CD4+ T cells (G) or the CD8^+^ T cells. (H) after G-CSF stimulation for 72 h *in vitro*. Error bars represent the mean ± SEM values from 3 independent healthy donors.

