## Supplemental Figures for "Three-dimensional genome structure and chromatin accessibility reorganization during *in vivo* induction of human T cell tolerance"

### Figure S1. Quality control for sequencing data.

A

| | Clean data | Reported_pairs(%) | Merged | Reported_pairs | Validate pairs | Validate (%) |
|---|---|---|---|---|---|---|
| HD1-CD4 Tss | 439,751,039 | 71.7 | 1,212,237,926 | 67.69 | 588,297,096 | 48.53% |
| HD2-CD4 Tss | 402,022,978 | 70.0 | | | | |
| HD3-CD4 Tss | 370,463,909 | 67.0 | | | | |
| HD1-CD4 Ttol | 408,148,478 | 64.9 | 1,192,855,550 | 57.21 | 466,302,176 | 39.09% |
| HD2-CD4 Ttol | 385,154,926 | 63.5 | | | | |
| HD3-CD4 Ttol | 399,552,146 | 50.0 | | | | |
| HD1-CD8 Tss | 425,713,372 | 70.4 | 1,046,971,086 | 56.36 | 404,017,844 | 38.59% |
| HD2-CD8 Tss | 318,611,093 | 69.8 | | | | |
| HD3-CD8 Tss | 302,646,621 | 46.7 | | | | |
| HD1-CD8 Ttol | 430,694,017 | 62.5 | 1,077,998,897 | 59.88 | 404,660,262 | 37.54% |
| HD2-CD8 Ttol | 313,338,650 | 64.2 | | | | |
| HD3-CD8 Ttol | 333,966,230 | 50.4 | | | | |

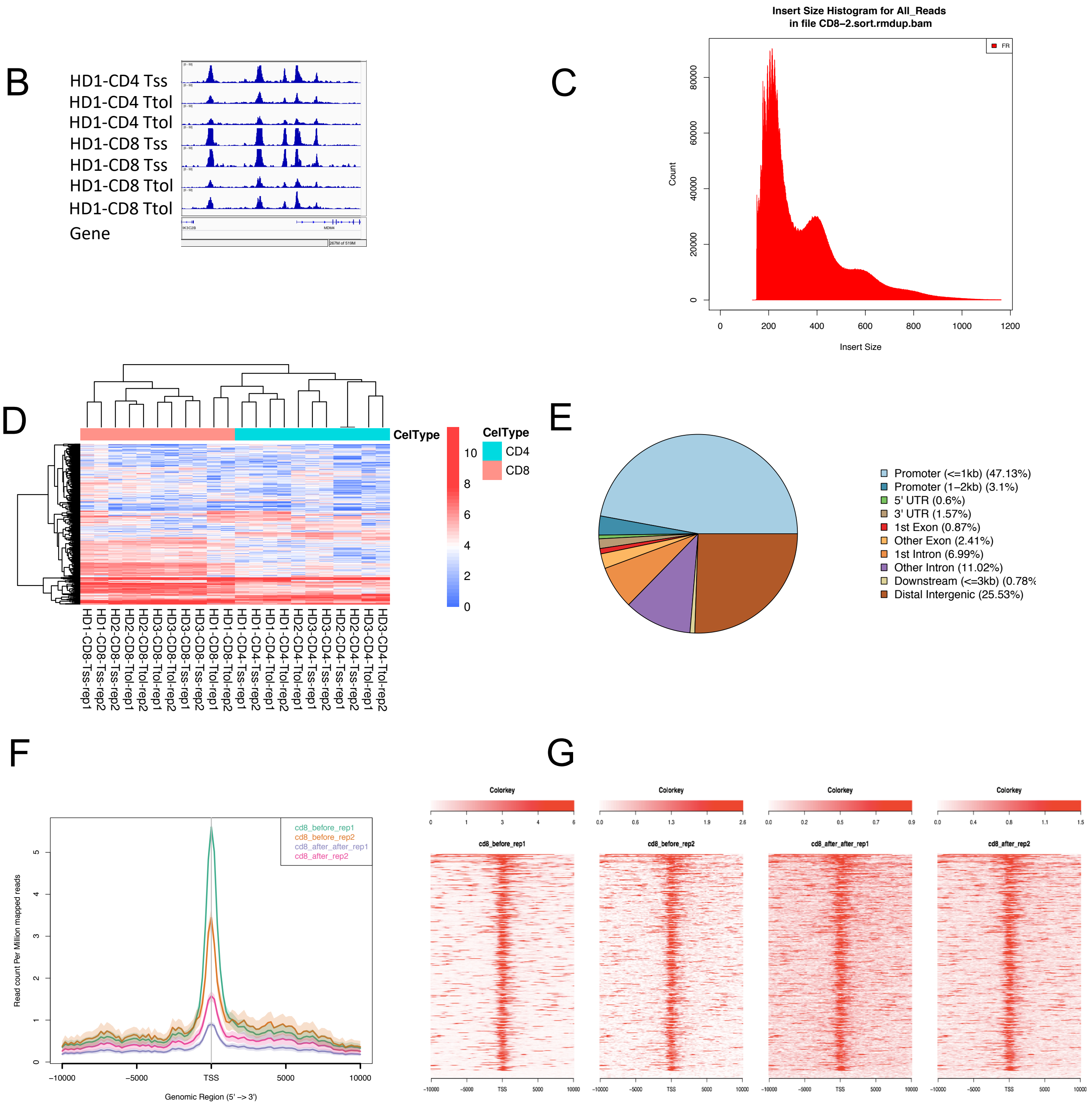

**Figure S2. Differentially expressed genes and differentially activated TFs in the CD4+ cells before and after mobilization.**

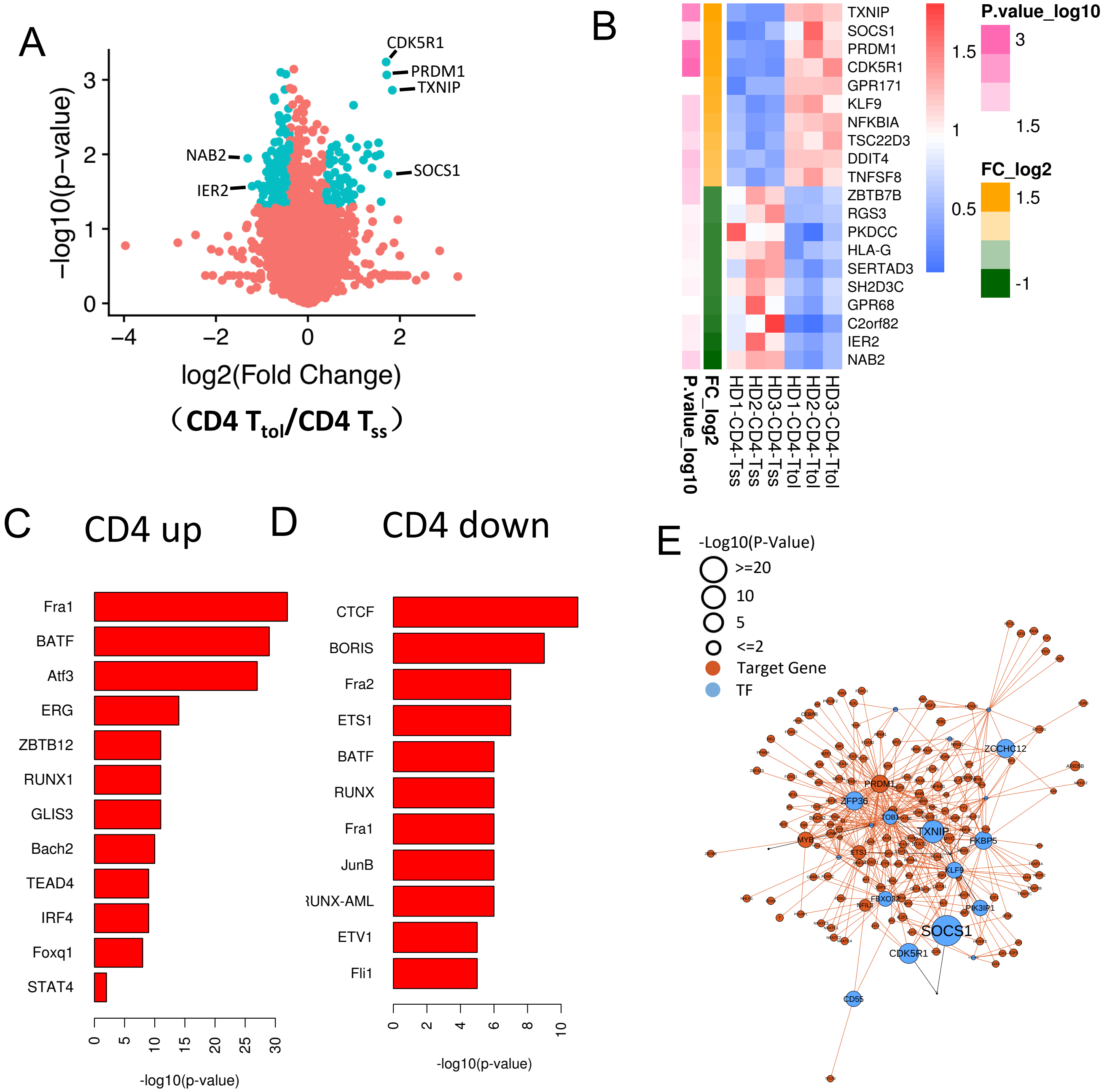

**Figure S3. Specific gene and chromatin structures in the CD4 and CD8 cells after mobilization.**

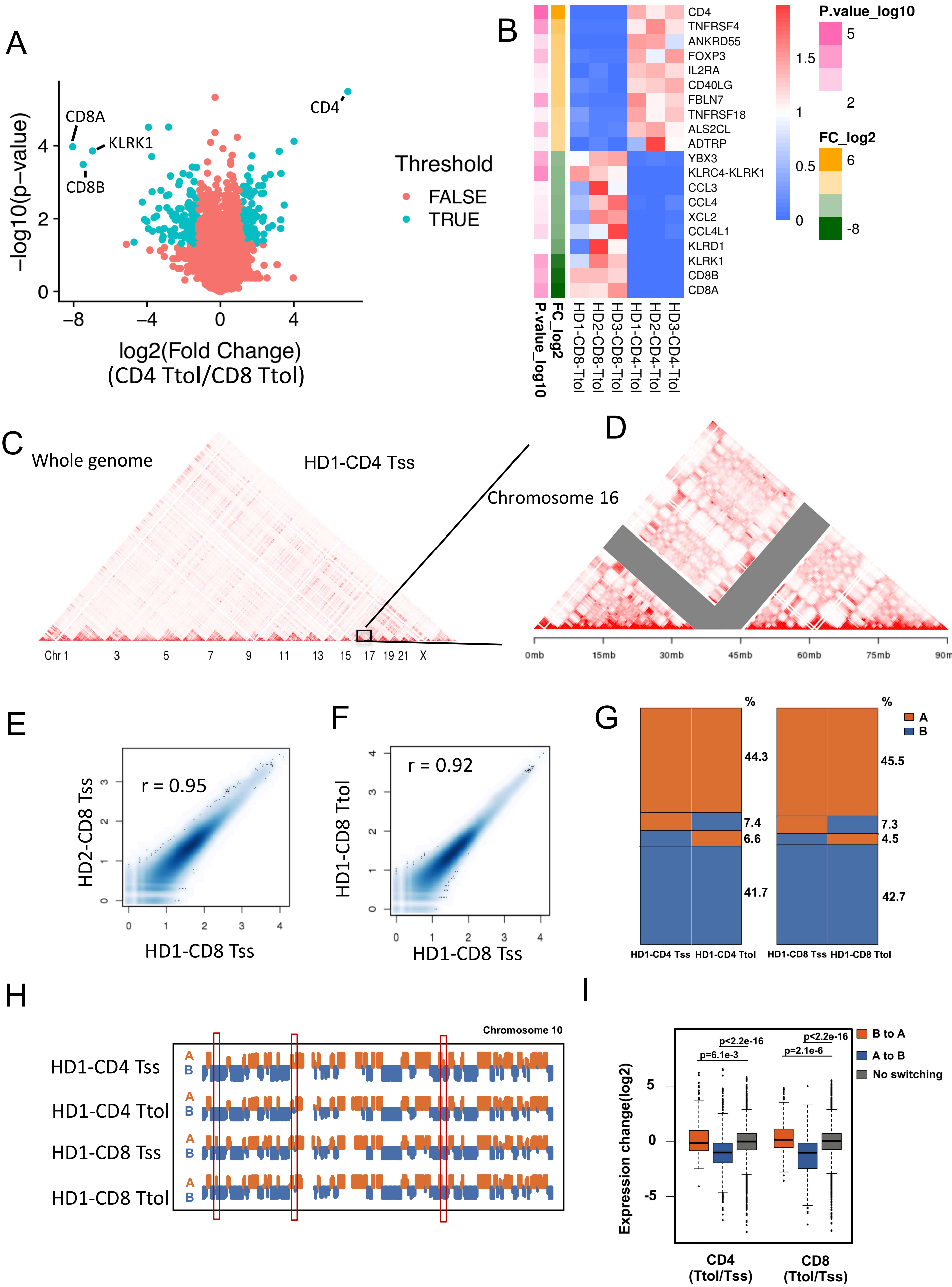

**Figure S4. TAD and loops structures are influenced by G-CSF.**

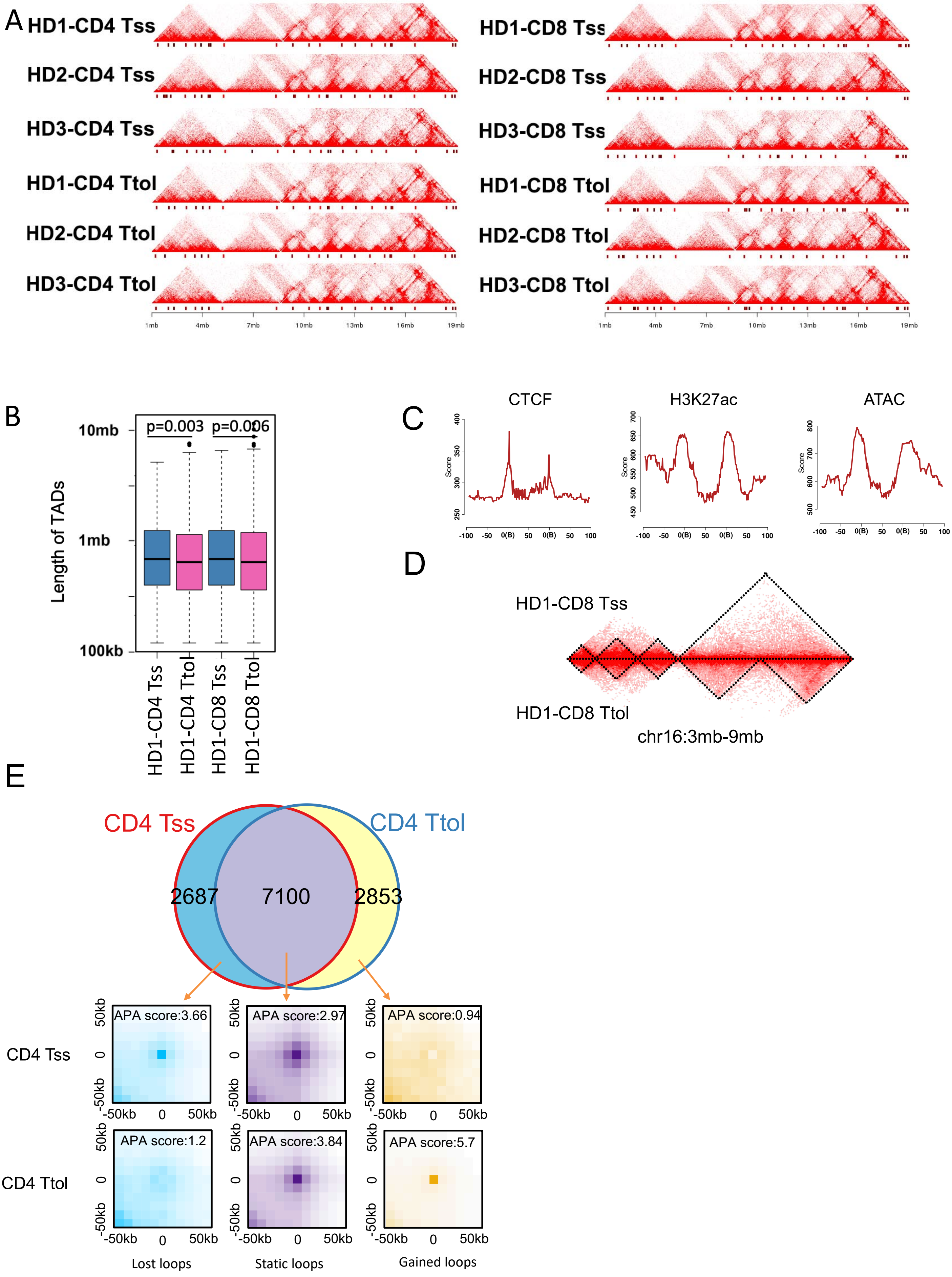

**Figure S5. STAT3 and CTCT are colocalized in the whole genome in the GM12878 and Jurkat cell lines.**

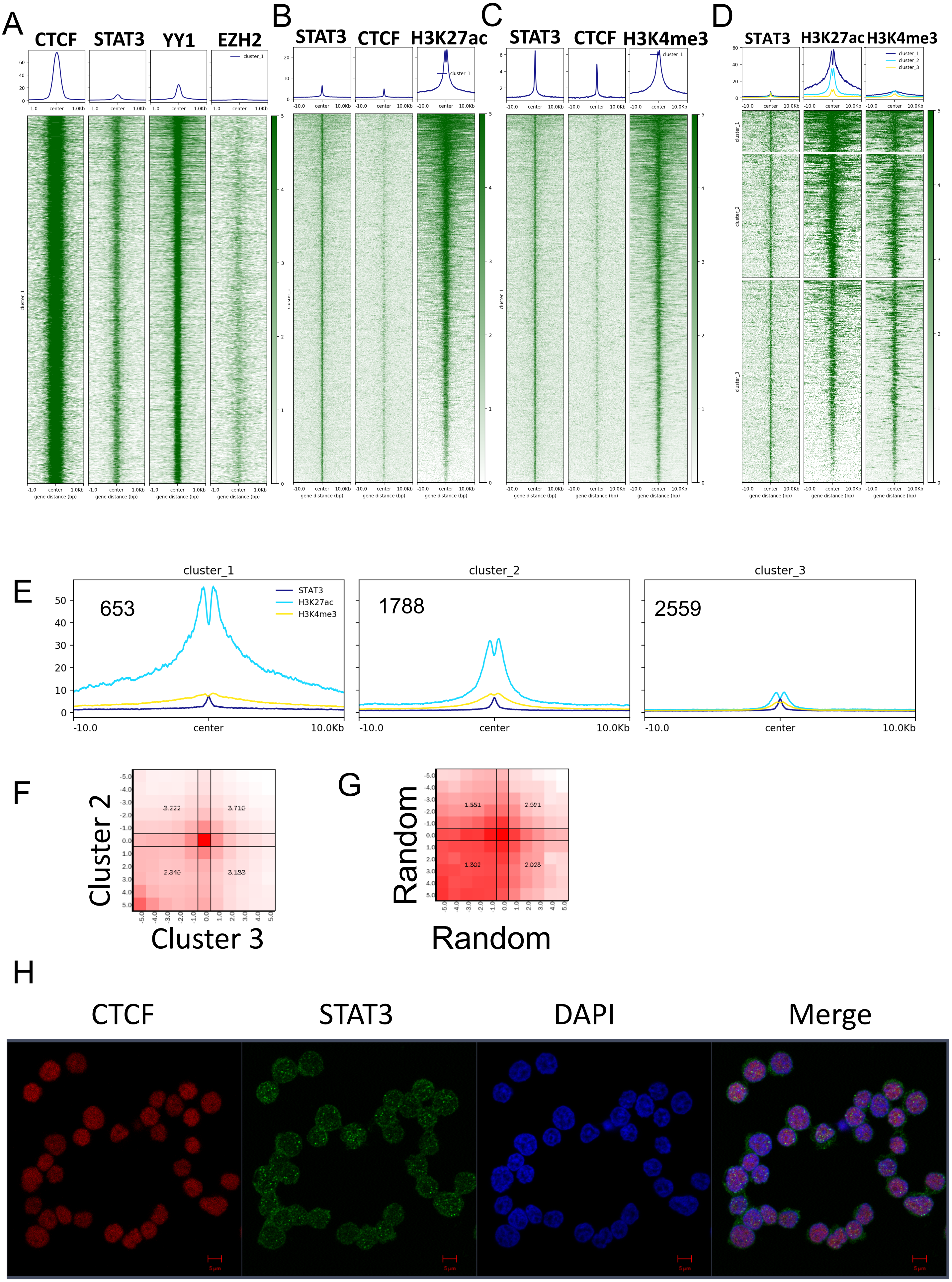

**Figure S6. High expression levels of SOCS1 promote exhaustion marker expression.**

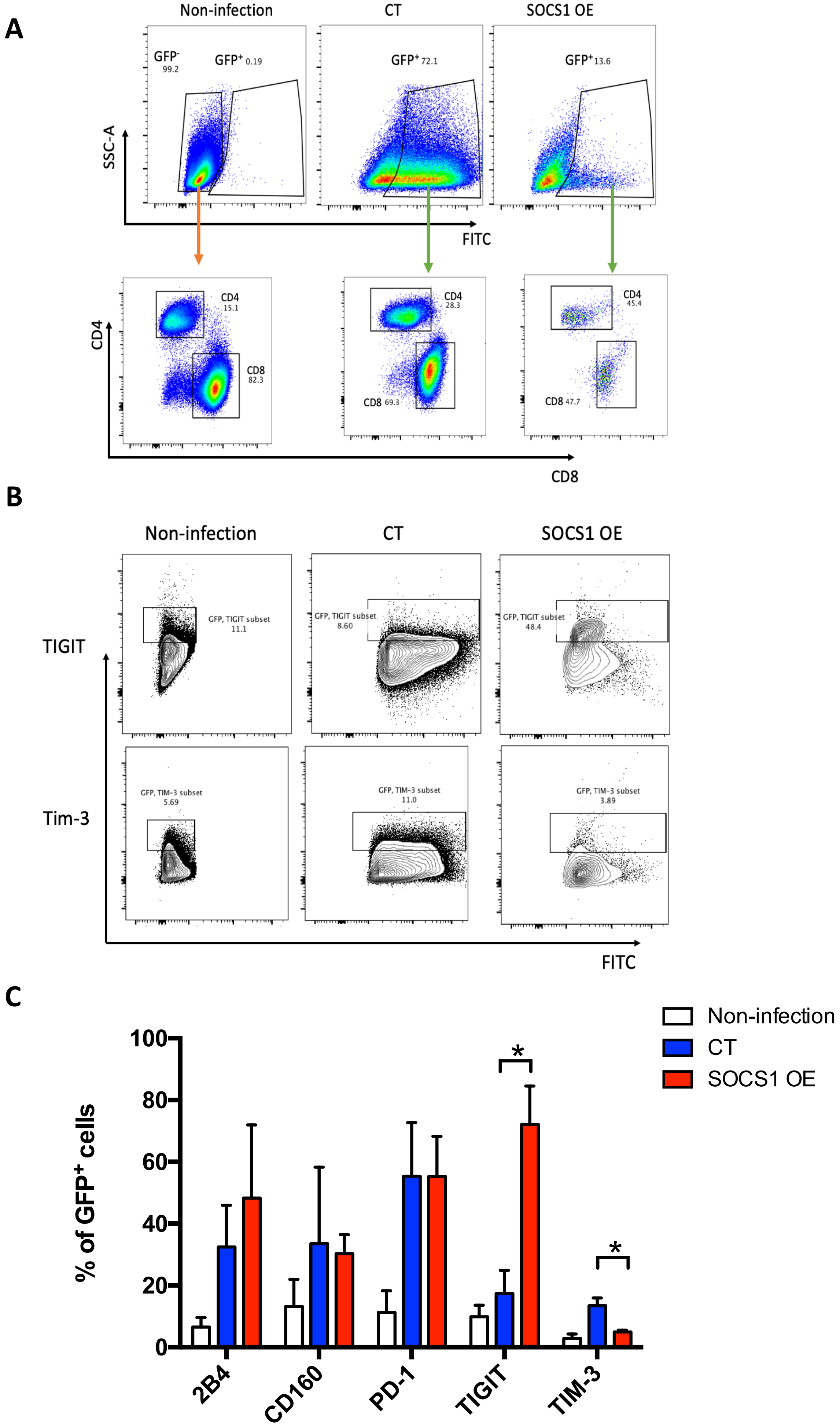

**FigureS7. Cytokine secretion level in T cells after SOCS1 overexpression *in vitro.***

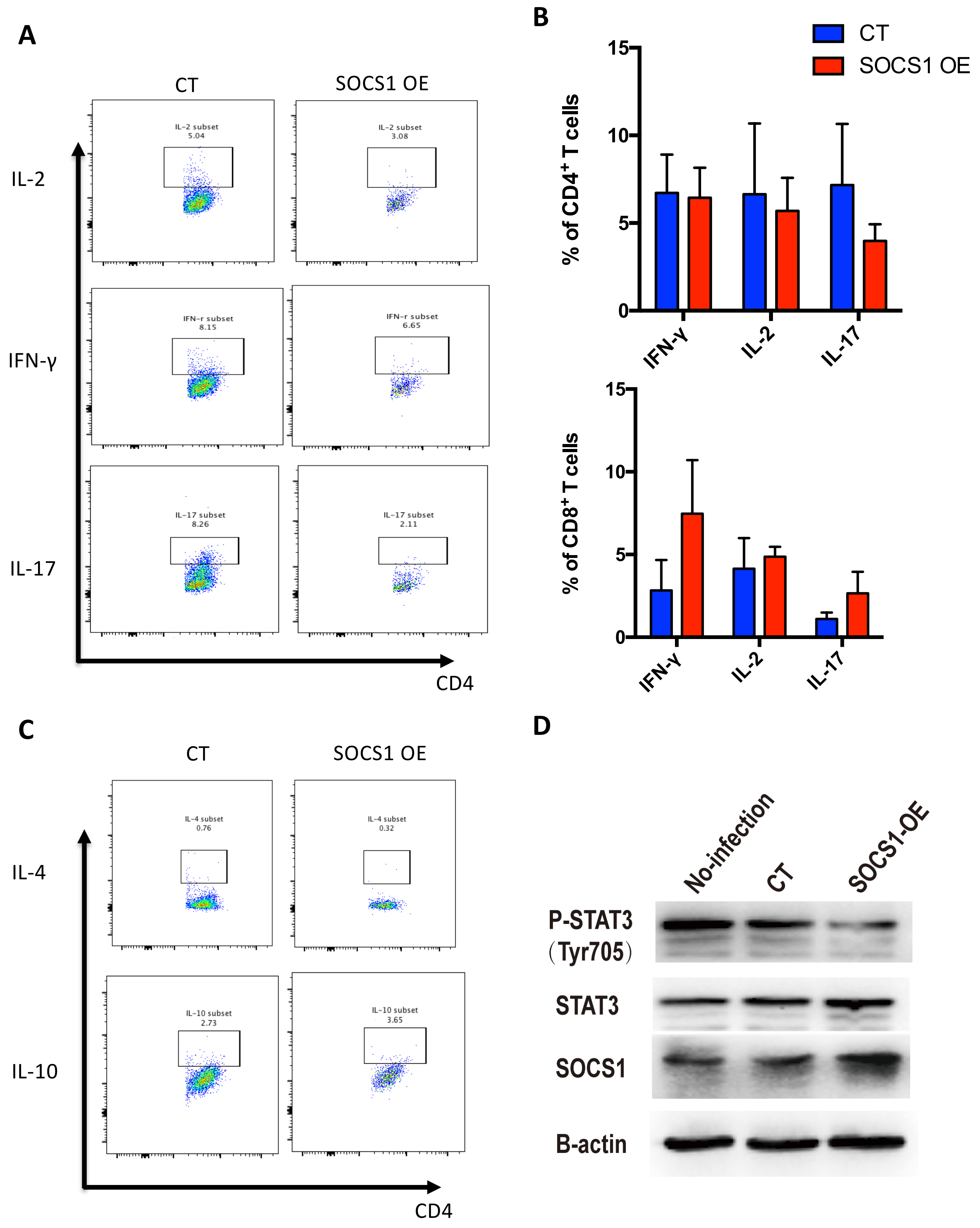

**FigureS8. IL-10 secretion in tolerant T cells *in vitro* after decreasing SOCS1 expression.**

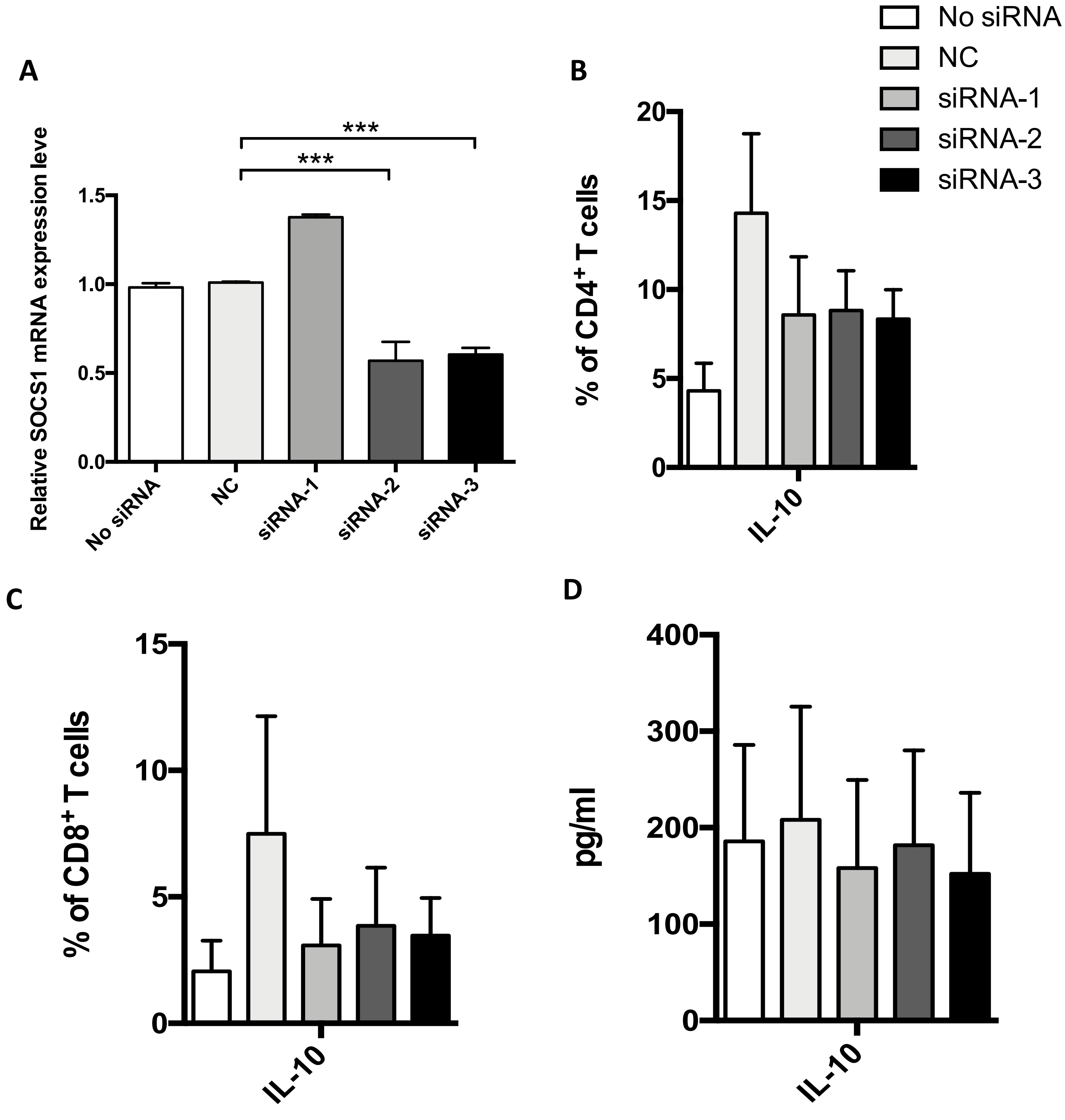

**Figure S9. G-CSF elevated the SOCS1 expression levels and suppressed the Th1 phenotype in T cells *in vitro***

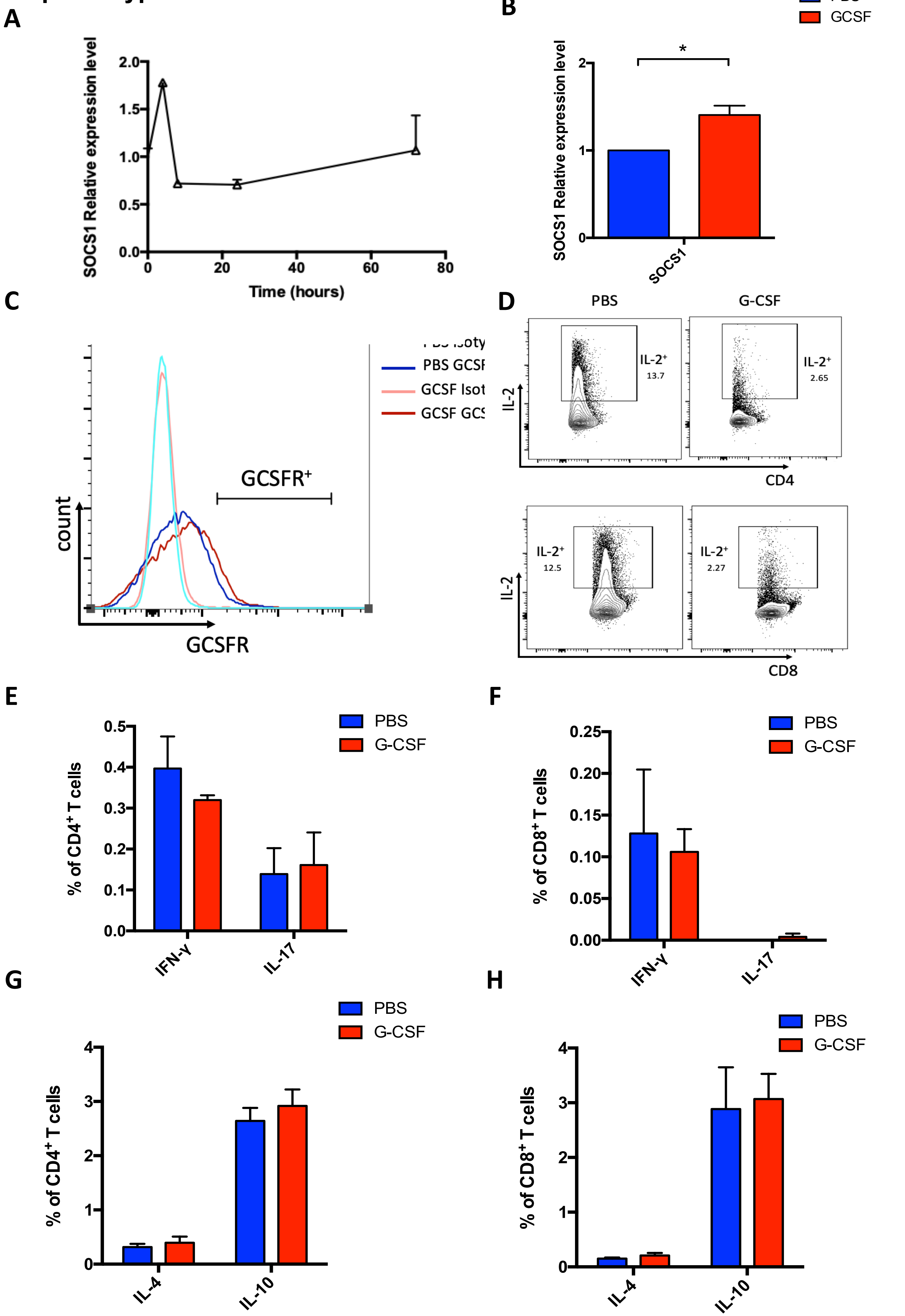
